# Processes essential for *Physcomitrium patens* protonemal development require distinct levels of total activity provided by functionally redundant PpROP GTPases

**DOI:** 10.1101/2024.12.20.629792

**Authors:** Aude Le Bail, Benedikt Kost, Janina Nüssel, Tamara Isabeau Lolis, David Koch, Hildegard Voll, Sylwia Schulmeister, Karin Ljung, Maria Ntefidou

## Abstract

RHO GTPases are key regulators of cellular and developmental processes in most eukaryotic organisms. ROPs (RHO of plants) constitute a plant-specific RHO subfamily. ROP families expanded and functionally diversified during the evolution of structurally complex vascular plants, but generally contain few members in non-vascular plants with ancient features. In vascular and non-vascular plants, ROP proteins are required for cell polarization, directional cell expansion, and mitotic cell plate positioning. All these processes are impaired by the disruption of PpROP activity in the non-vascular moss *Physcomitrium patens*. The aim of the study presented here was to further characterize PpROP functions during *P. patens* protonemal development by a) knocking out individually or in all possible combinations each of the four Rp*ROP* genes, which encode nearly identical proteins, b) complementing knock-out mutants with WT or mutant PpROP isoforms, or with heterologous homologs, and c) performing overexpression experiments. PpROPs were found to have additional previously unknown functions in the regulation of cell proliferation, caulonema differentiation, and gametophore formation. Furthermore, different cellular and developmental processes were shown to require distinct levels of total PpROP activity, rather than individual PpROP isoforms. Implications of the remarkable sequence conservation and functional integration within the PpROP protein family are discussed.

**One-sentence summary:** Knock-out, complementation, and overexpression experiments further defined PpROP functions in *P. patens* development, and demonstrated their dependence on distinct levels of total PpROP activity.

## Introduction

RHO (RAS homologous) family small GTPases (guanosine triphosphate hydrolases) are expressed in most eukaryotic organisms (Boureux et al., 2007) and play key roles in the regulation of essential cellular processes with important functions in the development of multicellular tissues and organs (Etienne-Manneville and Hall, 2002; Kawano et al., 2014). Most RHO GTPases are associated with the plasma membrane (PM) based on posttranslational prenylation, interact with different effectors to trigger downstream signaling specifically in the GTP-bound confirmation, and are inactive when bound to GDP after GTP hydrolysis. While regulatory factors that control RHO activity by promoting GDP for GTP exchange, by stimulating GTP hydrolysis or by modulating PM-association are closely related in different organisms (Berken et al., 2005; Kost, 2010; Hodge and Ridley, 2016), RHO-dependent downstream signaling appears to be much less conserved. Different families of RHO effectors are controlling similar cellular processes in animals and in plants (Heasman and Ridley, 2008; Yalovsky et al., 2008; Müller, 2023; Ntefidou et al., 2023). Furthermore, RHO effectors playing key roles in the regulation of important cellular processes in complex vascular plants are either entirely missing (Eklund et al., 2010; Ou and Yi, 2022) or have completely different functions in non-vascular plants with more ancient features (Ntefidou et al., 2023).

RHO functions in the control of developmentally relevant cellular processes including polarization, directional growth, motility, and division have been extensively characterized both in animals and in plants. However, the direct demonstration of RHO functions in tissue and organ development based on the investigation of loss-of-function mutants has been hampered by substantial redundancy within large families of RHO proteins typically expressed in complex multicellular organisms and, possibly, by essential roles of at least some of these proteins in the development of such organisms (Mulvey and Dolan, 2023b).

ROP (RHO of Plants) functions have been most extensively studied in *Arabidopsis thaliana*, a complex flowering plant with an elaborate vascular system, which expresses 11 AtROPs sharing 80-98 % amino acid sequence identity (Li et al., 1998; Winge et al., 2000). The functional characterization of these proteins largely depended on the investigation 1) of loss-of-function mutants, and 2) of effects of overexpressing either wild type (WT) AtROPs, or mutant versions of these proteins locked in the GTP-bound conformation (constitutively active) or displaying reduced nucleotide affinity (moderate reduction: fast-cycling, strong reduction: dominant negative). Different AtROPs were determined to play distinct but partially overlapping (Feiguelman et al., 2018) essential roles in the control of the following cellular processes: 1) extremely polarized tip growth displayed by pollen tubes (Li et al., 1999; Luo et al., 2017) and root hairs (Molendijk et al., 2001; Denninger et al., 2019), 2) diffuse directional expansion of leaf epidermal pavement cells (Fu et al., 2005; Lauster et al., 2022), 3) morphogenesis of single-celled trichomes (Liu et al., 2023), and 4) mitotic cell plate positioning required for tissue patterning in developing roots (Roszak et al., 2021).

Interestingly, Arabidopsis mutants disrupted in the expression of multiple At*ROP* genes show severely aberrant root hair and/or epidermal leaf pavement cells morphogenesis, but no substantial defects in tissue organization and organ development (Fu et al., 2005; Ren et al., 2016). Similarly, the development of the liverwort *Marchantia polymorpha* is not strongly affected by knocking out the single Mp*ROP* gene identified in this non-vascular plant, even though defects in the formation of selected tissues and organs were observed (Mulvey and Dolan, 2023b). The relatively weak phenotype of *M. polymorpha rop* knock-out mutants obviously cannot be attributed to genetic redundancy. ROP signaling therefore only appears to play a minor role in controlling the development in this liverwort, which displays ancient features presumably similar to those of extinct common ancestors of all land plants. By contrast, growth and development of the closely related non-vascular moss *P. patens* are severely disrupted in knock-out or knock-down mutants completely lacking or displaying minimal ROP activity (Burkart et al., 2015; Cheng et al., 2020; Yi and Goshima, 2020; Bao et al., 2022). These mutants only form tiny colonies of irregularly shaped cells, which are unable to directionally expand, differentiate or form organized tissues. The phenotype of these mutants established that *P. patens* development strictly depends on the total activity of the four nearly identical PpROPs expressed in this moss, which share 99.5-100 % amino acid identity and display sequence variability only outside of known functional domains at positions 148 and 149 (Eklund et al., 2010). However, currently available data concerning PpROP functions leave several important questions unanswered: 1) does PpROP activity only exert previously identified functions in the control or cell polarization, directional cell expansion and mitotic cell plate positioning, or is this activity also required for other cellular and developmental processes?, 2) are different processes controlled by individual ROP isoforms or by the combined activity of multiple PpROPs?, 3) are the four PpROPs functionally redundant or do they differentially contribute to total PpROP activity?, and 4) which evolutionary mechanisms may account for the origin and maintenance of the PpROP protein family with its exceptionally high amino acid sequence conservation?

The development of haploid *P. pates* gametophytes, which are composed of protonemal filaments and gametophores, is an ideal system to further characterize PpROP functions. This process 1) dominates the *P. patens* life cycle, 2) relies on cell polarization, tip growth, strictly controlled cell plate positioning, and additional cellular or developmental processes, and 3) is highly amenable to genetic manipulation based on homologous recombination and CRISPR/Cas (Schaefer and Zrÿd, 1997; Cove, 2005; Collonnier et al., 2017; Rensing et al., 2020). Gametophyte development starts with the germination of haploid spores, which results in the formation of branched filamentous protonemata. These structures can also be induced to develop *in vitro* from isolated protonemal protoplasts or explants, with protoplast-derived colonies more closely recapitulating normal development (Grimsley et al., 1977; Kofuji and Hasebe, 2014). A single apical initial cell at the tip of each protonemal filament expands by tip growth and regularly divides transversely, resulting in filament elongation. The association of different PpROPs with the PM of apical initial cells specifically at the tip, as well as underneath the transversal cell wall separating these cells from subapical cells, is consistent with essential functions of PpROP activity in tip growth and cell plate positing (Le Bail et al., 2019; Cheng et al., 2020; Yi and Goshima, 2020). While young protonemal filaments display chloronemal characteristics, the initial cells at their tips undergo gradual caulonema differentiation, which is associated with 1) an increased rate of tip growth and proliferation, 2) a reduction in chloroplast size and number, and 3) the formation of oblique rather than perpendicular transversal cell walls (Jaeger and Moody, 2021). Interestingly, caulonema differentiation is stimulated by the phytohormone auxin (Jang and Dolan, 2011) and has recently been demonstrated to be effectively blocked by the RpROP effector PpRIC downstream of auxin-induced changes in gene expression (Ntefidou et al., 2023). Filament branching as well as the formation of lateral buds developing into gametophores are initiated by the asymmetric division of subapical caulonemal cells. As this process is defective in mutants lacking 3 of the four PpROPs, it also appears to be controlled by PpROP activity (Yi and Goshima, 2020). Mature gametophores are composed of leafy shoots with leaf-like phyllids and filamentous rhizoids, which are free of chloroplasts but otherwise closely resemble caulonemal filaments and elongate based on the same mechanisms. Reproductive organs produced at the tip of mature gametophores mediate sexual reproduction, which initiates sporophyte development that corresponds to the short diploid phase of the life cycle. To address open questions concerning PpROP functions during different phases of *P. patens* gametophyte development, systematic knock-out, complementation and overexpression experiments were performed.

## Results

### Individual Pp*ROP* genes are not essential for protonemal development

Previously reported quantitative polymerase chain reaction (RT-qPCR) and RNA-sequencing data demonstrated similar expression levels of all four Pp*ROP* genes in developing protonemata (Perroud et al., 2018, Ntefidou et al., 2023). RT-qPCR analyses performed here essentially confirmed these observations, although Pp*ROP4* was found to display about 2x higher transcript levels as compared to the other three genes (Supplementary Fig. S1). Based on these data, all four PpROPs appear to contribute to the control of protonemal growth and/or differentiation.

To systematically investigate PpROP functions in protonemal development, homologous recombination (Schaefer et al., 1991) was employed to replace all Pp*ROP* genes individually and in every possible combination by antibiotic resistance genes. Knockout (KO) lines obtained were verified by PCR genotyping (Supplementary Figs. S2A and B, S3 and S4). Higher order mutants were generated sequentially, starting with *rop1*, *rop2*, *rop3*, and *rop4* single KO lines (Supplementary Fig. S2C, upper panel). To enable comparative quantitative characterization of KO phenotypes, protonemata regenerated from isolated protoplasts were investigated at defined developmental stages. At least two independent lines per genotype were generally characterized, except for the *rop2*, *rop4*, r*op4/1* and *rop2/4/1* genotypes, which were only obtained once. Independently generated genotypically identical lines in all cases displayed indistinguishable phenotypes.

None of the four single *rop* KO mutants (*rop*^1xKO^) displayed detectable defects in a) the length of fully expanded subapical chloronemal or caulonemal cells (Supplementary Fig. S5A), b) the morphology of the tips of chloronemal or caulonemal filaments (Supplementary Fig. S5B), c) the rate of caulonema differentiation (Supplementary Fig. S5C), which is enhanced in a KO mutant lacking the PpROP effector PpRIC (*ric-1*^KO^, Ntefidou et al., 2023), or d) the size or morphology of 5-day-old protonemata or of 5 week-old colonies (Supplementary Fig. S5D-F). These observations establish that each individual *ROP* gene is dispensable for cell expansion and differentiation during protonemal development.

### Mutants lacking any combination of two Pp*ROP* genes display enhanced caulonema differentiation

The six *rop*^2xKO^ mutants lacking two Pp*ROP* genes in all possible combinations did not show detectable defects in cell expansion or morphology in 5-day-old protonemata, except for an apparent minor decrease in the size of *rop4/1* protonemata (Fig. 1A, B, D and F). However, at the same developmental stage caulonema differentiation was substantially enhanced to a similar extent in all *rop*^2xKO^ mutants (Fig. 1C), establishing an essential function of PpROP activity in the inhibition of this process. Caulonema differentiation was proposed to be triggered when elongating apical chloronemal cells reach a threshold length (Jang and Dolan 2011). Since the length of chloronemal cells was not affected in *rop*^2xKO^ mutants (Fig. 1A), enhanced caulonema differentiation displayed by these mutants is not a secondary effect of altered cell expansion, but appears to be a direct consequence of reduced PpROP activity.

**Figure 1.**
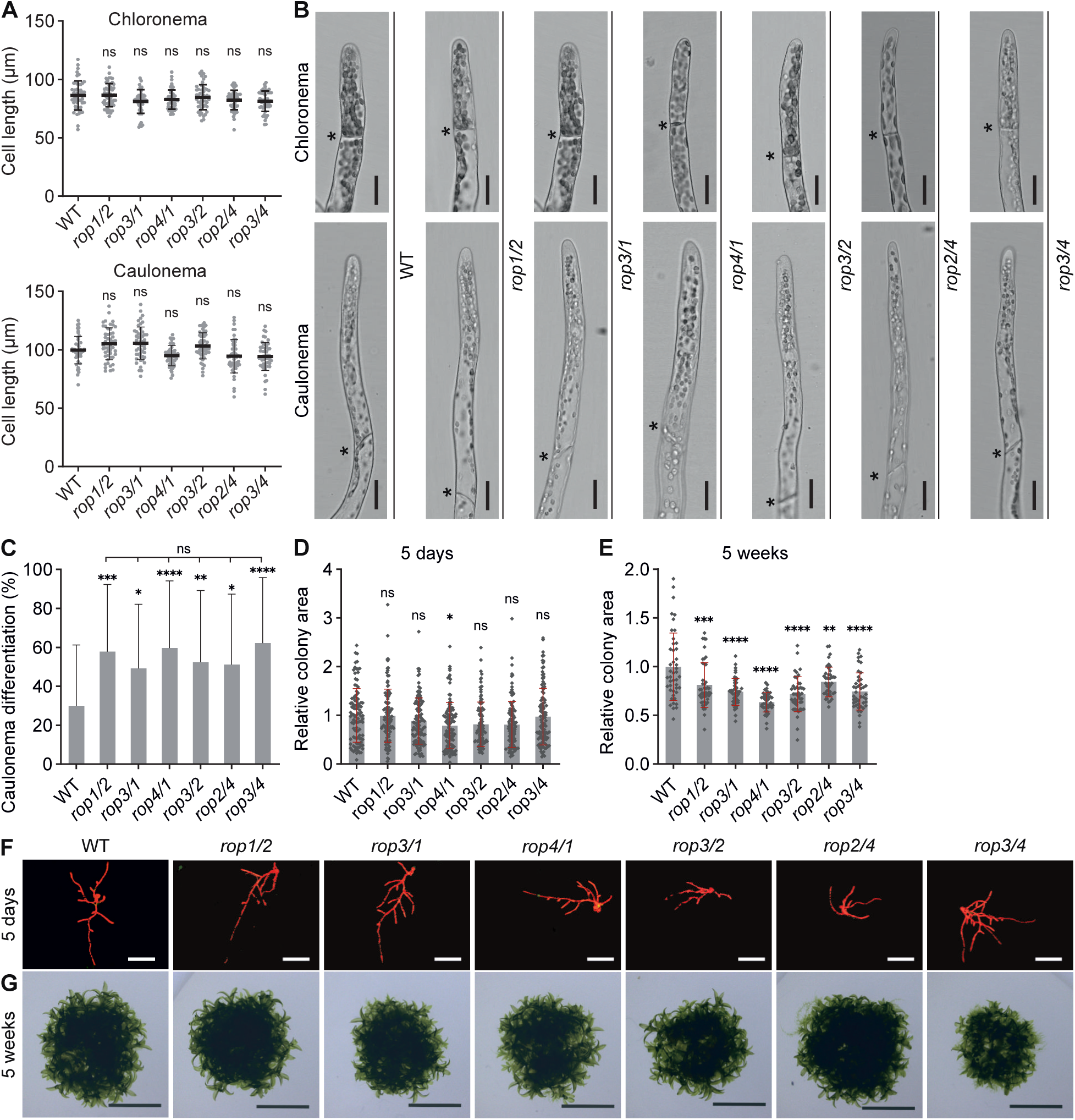
Knockout of two Pp*ROP*s (*rop*^2xKO^) promotes caulonema differentiation. Graphs and images based on 5-day-old protonemata or 5-week-old colonies regenerated from protoplasts with the indicated genotypes of double Pp*ROP* knockout mutants and WT using culture conditions listed in Supplementary Table S4. **A)** Average subapical cell length of chloronema and caulonema cells in 5-day-old protonemata. n = 50 cells were analyzed per genotype. The experiment was performed three times with consistent results. **B)** Bright field images of 5-day-old chloronemal and caulonemal filament tips. Asterisks indicate the cell wall between neighboring cells. Scale bars: 25 µm. **C)** Average percentage of caulonema differentiation in 5-day-old protonemal filaments with at least three cells as determined by microscopic observation. n = 60 colonies per genotype were measured in 3 independent experiments. **D-G)** Average size (area) of 5-day-old protonemata **(D)** or 5-week-old colonies **(E)** as determined based on microscopic imaging of chlorophyll autofluorescence **(F**, 5-day-old) or pictures recorded using a stereo microscope **(G**, 5-week-old). The mean value of WT was used as a calibrator (relative area = 1). n = 105 colonies per genotype were measured in 3 independent experiments (**D**), or n = 42 colonies per genotype were measured. The experiment was performed three times with consistent results (**E**). Scale bars: 400 µm **(F)**, 10 mm **(G)**. (**A, C-E)** Error bars: standard deviation (SD), dots represent individual data points. Statistical analysis by one-way ANOVA/Tukey’s test (pairwise comparisons to WT and relevant pairwise comparisons are displayed, all others see Supplementary Data Set S3): ^ns^ *P* > 0.05 (not significant), * *P* ≤ 0.05, ** *P* ≤ 0.01, *** *P* ≤ 0.001, **** *P* ≤ 0.0001.

As none of the *rop*^1xKO^ mutants showed enhanced caulonema differentiation (Supplementary Fig. S5C), any combination of three Pp*ROP* genes evidently is sufficient for the inhibition of this process in normally developing protonemata, which has previously been shown to also require the PpROP effector PpRIC (Ntefidou et al., 2023). PpRIC was demonstrated to inhibit caulonema differentiation downstream of gene expression controlled by indole-3-acetic acid (IAA), an auxin-family phytohormone with a key role in the stimulation of this process (Jaeger and Moody, 2021). To further investigate whether PpROP activity and PpRIC act together in the same pathway to inhibit caulonema differentiation as proposed by Ntefidou et al. (2023), IAA levels, a marker for auxin transport and auxin dependent-gene expression were analyzed in *rop*^2xKO^ mutants. Like *ric-1*^KO^ mutants (Ntefidou et al., 2023), all *rop*^2xKO^ mutants were found a) to contain wild-type IAA levels (Supplementary Fig. S6A), b) to display normal accumulation of the key auxin transporter PpPINA (PIN-FORMED A auxin efflux carrier; Viaene et al., 2014) specifically in the PM at the tip of apical protonemal cells (Supplementary Fig. S6B), and c) to express at normal levels different auxin-responsive genes (Supplementary Fig. S6C), which are required for auxin-induced caulonema differentiation (Pp*RSL1*; *ROOTHAIR DEFECTIVE SIX-LIKE1*; Jang et al., 2011), or have essential functions in auxin biosynthesis (Pp*SHI1*; *SHORT INTERNODES1*; Eklund et al., 2010b), transport (Pp*PINA*) or signaling (Pp*AUX1*; *AUXIN RESISTANT 1*; Lavy et al., 2016). These observations establish that PpROP activity, like PpRIC, inhibits caulonema differentiation without affecting auxin-controlled gene expression, supporting the notion that PpROP activity regulates this process via PpRIC stimulation.

Consistent with a considerably higher growth rate of caulonemal as compared to chloronemal filaments (Cove and Knight, 1993), enhanced caulonema differentiation substantially enlarged the size of 5-week-old *ric-1*^KO^ colonies (Ntefidou et al., 2023). By contrast, despite enhanced caulonema differentiation, the size of 5-week-old *rop*^2xKO^ colonies was significantly reduced (Fig. 1E and G), although as discussed above, 5-day-old *rop*^2xKO^ protonemata did not display substantial growth defects (Fig. 1A, B, D and F). Based on these observations, PpRIC appears to be exclusively required for the inhibition of caulonema differentiation, whereas RpROP activity, as previously demonstrated, also promotes cell and protonemata expansion, evidently by stimulating other effectors. However, it is important to note that *rop*^2xKO^ protonemata displayed minimal growth defects, which only became apparent late during development and did not prevent the detection of enhanced caulonema differentiation at an earlier stage.

### Cell and protonemal expansion are severely disrupted in mutants lacking any combination of three Pp*ROP* genes

Consistent with a previously reported phenotypic analysis of *P. patens rop1/2/3* and *rop2/3/4* triple KO mutants (Yi and Goshima, 2020), 5-day-old protonemata of each of the four *rop*^3xKO^ mutants lacking three Pp*ROP* genes in all possible combinations displayed severe growth defects (Fig. 2A, B, D and G). Already at this early developmental stage, the length of apical and subapical chloronemal and caulonemal cells (Fig. 2A and B), as well as the size of protonemata (Fig. 2D and G), were substantially decreased. Interestingly, at the same developmental stage and by contrast to *rop*^2xKO^ mutants, *rop*^3xKO^ mutants did not display enhanced caulonema differentiation (Fig. 2C). As caulonema differentiation seems to depend on a cell length threshold (Jang and Dolan, 2011), reduced cell expansion appears to inhibit this process in *rop*^3xKO^ protonemata.

**Figure 2.**
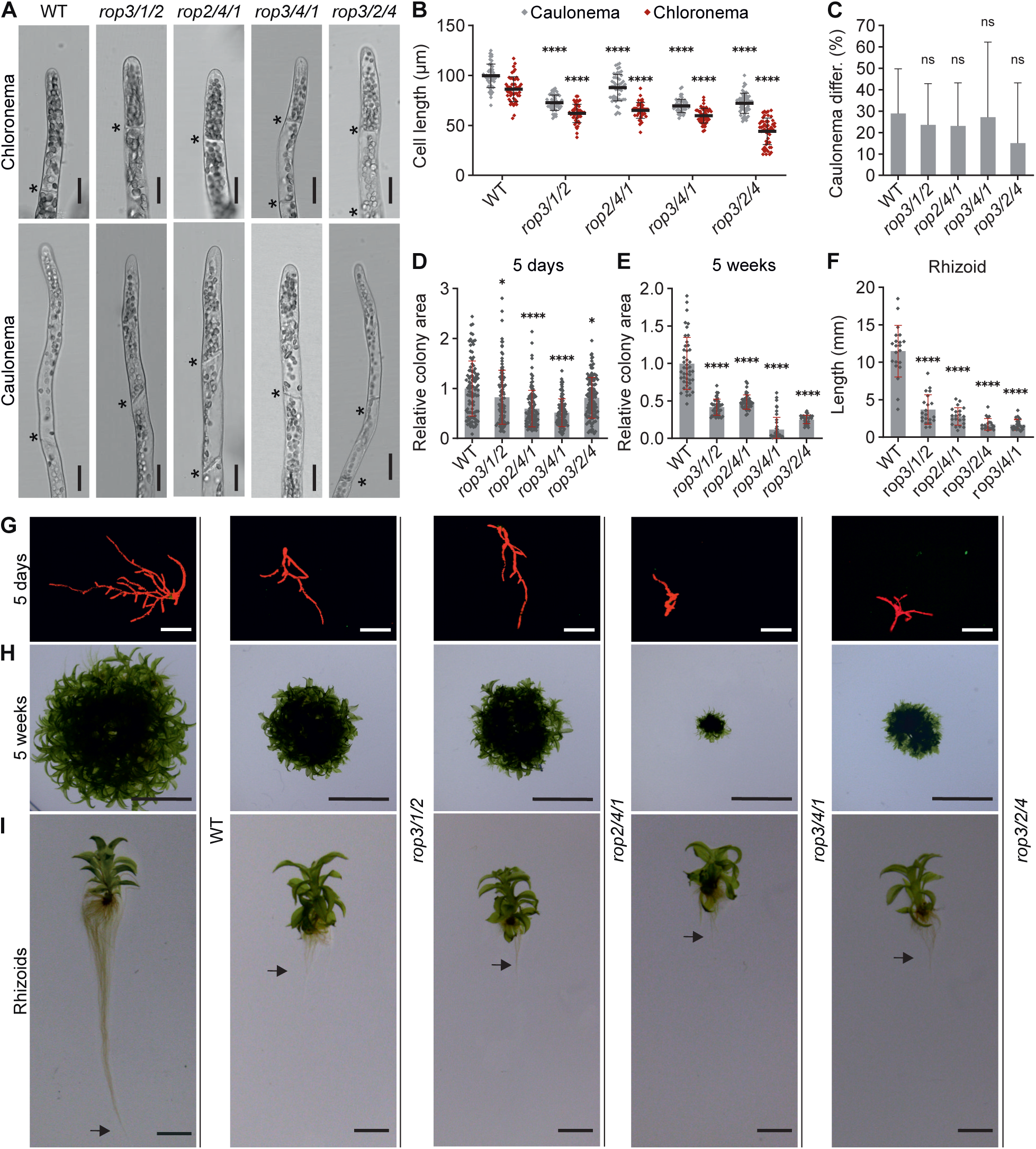
Knockout of three Pp*ROP*s (*rop*^3xKO^) inhibits tip growth of protonemata and rhizoids. Graphs and images based on 5-day-old protonemata or 5-week-old colonies regenerated from protoplasts with the indicated genotypes of triple Pp*ROP* knockout mutants and WT cultivated on media listed in Supplementary Table S4. **A)** Bright field micrographs of 5-day-old chloronemal and caulonemal filament tips. Asterisks indicate the cell wall between neighboring cells. Scale bars: 25 µm. **B)** Average subapical cell length of chloronemal and caulonemal cells in 5-day-old protonemata. n = 50 cells per genotype were measured. The experiment was repeated three times with consistent results. **C)** Average percentage of caulonema differentiation in 5-day-old protonemal filaments with at least three cells as determined by microscopic observation. n = 60 colonies per genotype were measured in 3 independent experiments. **D, E, G, H)** Average size (area) of 5-day-old protonemata **(D)** or 5-week-old colonies **(E)** as determined based on microscopic imaging of chlorophyll autofluorescence **(G)** or pictures recorded using a stereo microscope **(H)**. Scale bars: 400 µm **(G)**, 10 mm **(H)**. The mean value of WT was used as a calibrator (relative area = 1). n = 105 colonies per genotype were measured in 3 independent experiments **(D),** or n = 45 colonies per genotype were measured. The experiment was repeated three times with consistent results **(E)**. **F** and **I)** Average length of rhizoids of 5-week-old gametophores **(F)** based on pictures recorded using a stereo microscope (arrows indicate the end of rhizoids) **(I)**. Scale bars: 5 mm. n = 21 rhizoids per genotype were measured. The experiment was repeated three times with consistent results. **B-F)** Error bars: standard deviation (SD), dots represent individual data points. Statistical analysis by two-way ANOVA/Tukey’s test **(B)** or one-way ANOVA/Tukey’s test **(C-F)** (comparisons to WT are displayed, all others see Supplementary Data Set S3): ^ns^ *P* > 0.05 (not significant), * *P* ≤ 0.05, **** *P* ≤ 0.0001.

The analysis of 5-week-old *rop*^3xKO^ colonies revealed even more pronounced growth defects (Fig. 2E and H). Interestingly and consistent with previously reported observations (Yi and Goshima, 2020), at this developmental stage *rop*^3xKO^ colonies formed gametophores with small but morphologically essentially normal leafy shoots and even more severely stunted rhizoids (Fig. 2F, H and I), which like protonemata are composed of filaments that elongate based on apical tip growth. The phenotypic characterization of *rop*^3xKO^ protonemata and gametophores therefore not only confirms essential functions of PpROP activity in the control of tip growth but also suggests less important roles of this activity in other forms of directional cell expansion, which are required for leafy shoot development. Whereas tip growth is strongly reduced in all *rop*^3xKO^ mutants (Fig. 2), this process is only marginally affected in all *rop*^2xKO^ mutants (Fig. 1). The expression of any combination of two PpROPs therefore appears to be sufficient to support nearly normal tip growth.

### All four PpROPs are equally capable of promoting tip growth

Amino acid sequence identities of at least 99.5 % within the PpROP protein family, together with highly similar phenotypes of all *rop*^2xKO^ (Fig. 1) and *rop*^3xKO^ (Fig. 2) mutants revealed by quantitative characterization, suggest essentially equivalent functions in all four PpROPs in developing protonemata. However, cell and protonema expansion was reduced to a remarkably variable extent in *rop*^3xKO^ mutants expressing distinct single PpROP family members (Fig. 2B, D and E), indicating that these proteins may differ in their ability to promote tip growth. Alternatively, the observed phenotypic variability of *rop*^3xKO^ mutants may be caused a) by differential up-or down-regulation of the expression of the remaining Pp*ROP* gene after the disruption of the other members of the Pp*ROP* gene family, b) by different genetic backgrounds resulting from multiple rounds of gene replacement based on homologous recombination, and/or c) by the expression of variable sets of selectable marker genes resulting from this procedure.

RT-qPCR analysis established that in all four *rop*^3xKO^ mutants the remaining Pp*ROP* gene was expressed at the same level as the corresponding gene in the wild type background (Fig. 3A). This demonstrates that the phenotypic variability displayed by *rop*^3xKO^ mutants is not caused by differential feed-back control of transcript levels within the Pp*ROP* gene family. To distinguish between the other possible causes of phenotypic variability displayed by *rop*^3xKO^ mutants, the activity of all PpROP family members was directly compared by individually expressing each of them under the control of the same promoter in an identical genetic background. To this end, based on homologous recombination the coding sequence (exons and introns) and 3′ untranslated region (UTR) of the endogenous Pp*ROP1* gene in the *rop3/2/4* background was replaced by CDS (coding sequence) fragments encoding PpROP1, PpROP2, or PpROP3, which were attached to the Pp*ROP1*, Pp*ROP2*, or Pp*ROP3* 3′ UTR, respectively (Supplementary Fig. S2A and C: lower panel). As PpROP1 and PpROP4 share identical amino acid sequences, the Pp*ROP4* CDS was not included in this analysis. Resulting *rop*^4xKO^/*ROP1*^pro^:*ROPX* lines (at least two per genotype) were confirmed by PCR genotyping (Supplementary Fig. S2B) and in essence represent *rop*^4xKO^ mutants sharing an identical genetic background, which are complemented by the expression of different single PpROPs under the control of the endogenous Pp*ROP1* promoter. RT-qPCR analysis confirmed that each of the *ROP1*^pro^:*ROPX* genes was expressed in the *rop*^4xKO^ background at a similar level as the endogenous Pp*ROP1* gene in the *rop3/2/4* mutant or in WT protonema (Fig. 3B). Size and morphology of 5-day-old protonemata as well as of 5-week-old colonies formed by all *rop*^4xKO^/*ROP1*^pro^:*ROPX* lines were found to be indistinguishable (Fig. 3C-E), establishing that all four 4 PpROPs are equally capable of promoting tip growth.

**Figure 3.**
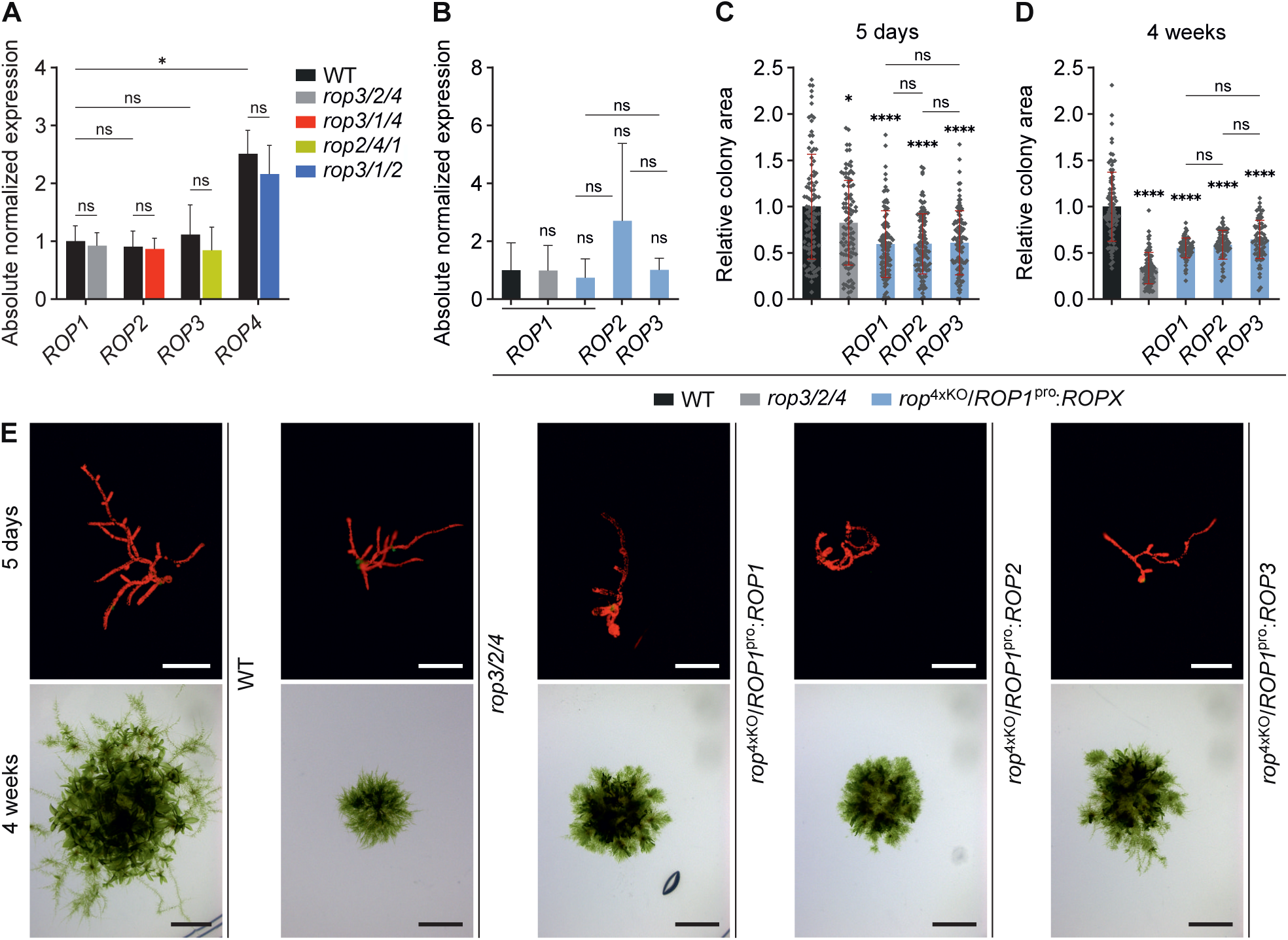
Pp*ROP*s are not altered in their gene expression in *rop*^3xKO^ mutants and redundantly complement the *rop*^4xKO^ mutant. **A** and **B)** qRT-qPCR of expression levels of Pp*ROP*s in 1-week-old protonemata cultivated through homogenization (Supplementary Table S4) with the indicated genotype: *rop*^3xKO^ mutants **(A)**, *rop*^4xKO^ mutant complemented with the coding and 3’ UTR sequence of Pp*ROP1* (Pp*ROP4* identical amino acid sequence) or Pp*ROP2* or Pp*ROP3* downstream of the Pp*ROP1* promoter (*rop*^4xKO^/*ROP1*^pro^:*ROPX*) **(B)** were compared to WT. Absolute transcript levels were determined based on standard curves using the value obtained for one WT replicate of Pp*ROP1* as a calibrator (relative expression = 1). Bars: mean of three biological and two technical replicates, error bars: standard error of the mean. The experiment was performed two times with consistent results. **C-E)** Average size (area) of 5-day-old protonemata **(C)** or 4-week-old colonies **(D)** regenerated from protoplasts with the indicated genotypes: i) WT, ii) *rop3/2/4*, iii) *rop*^4xKO^/*ROP1*^pro^:*ROPX* as determined based on microscopic imaging of chlorophyll autofluorescence **(E,** 5 days) or pictures recorded using a stereo microscope **(E,** 4 weeks). The mean value of WT was used as a calibrator (relative area = 1). n = 100 **(C)** or n = 80 **(D)** colonies per genotype were measured in 3 independent experiments. Error bars: standard deviation (SD), dots represent individual data points. Scale bars: 250 µm (**E** upper row, 5 days), 5 mm (**E** lower row, 4 weeks). Statistical analysis by two-way ANOVA/Bonferroni’s test **(A)** or one-way ANOVA/Tukey’s test **(B**-**D)** (pairwise comparisons to WT and relevant pairwise comparisons are displayed, all others provided in Supplementary Data Set S3): ^ns^ *P* > 0.05 (not significant), * *P* ≤ 0.05, **** *P* ≤ 0.0001.

### Disruption of all four Pp*ROP* genes completely abolishes polarized cell expansion and affects cell division

Two independent *rop*^4xKO^ lines were generated (Supplementary Fig. S2C: upper panel) by knocking out based on homologous recombination either Pp*ROP1* in the *rop3/2/4* background (*rop3/2/4/1*; Supplementary Fig. S2A and B) or Pp*ROP2* in the *rop3/4/1* background (*rop3/4/1/2*; Supplementary Fig. S3A and B). Both *rop*^4xKO^ lines displayed the same phenotype (Fig. 4A) as previously described *P. patens* mutants, in which the expression of all 4 *PpROP* genes was disrupted either by RNAi/microRNA-mediated downregulation (Burkart et al., 2015; Bao et al., 2022), by CRISPR/Cas9 (clustered regularly interspaced short palindromic repeats/CRISPR-associated nuclease 9)-mediated KO (Cheng et al., 2020), or by a combination of these approaches (Yi and Goshima, 2020). Mutants entirely or nearly completely lacking PpROP activity form tiny colonies of irregularly shaped cells (Fig. 4A). By contrast to cells containing at least one functional PpROP gene (*rop*^3xKO^), protonemal *rop*^4xKO^ cells were unable a) to undergo normal polarization, b) to maintain parallel orientation of cell division planes required for the development of linear filaments, or c) to directionally expand (Fig. 4A and B). When cultured for prolonged periods of time, *rop*^4xKO^ colonies continued to slowly grow based on cell division but failed to differentiate and did not develop gametophores. Interestingly, *rop*^4xKO^ lines essentially phenocopy *P. patens* KO mutants lacking geranylgeranyltransferase type I activity (Thole et al., 2014), suggesting that PpROPs require prenylation by this activity to be functional (Bao et al., 2022).

**Figure 4.**
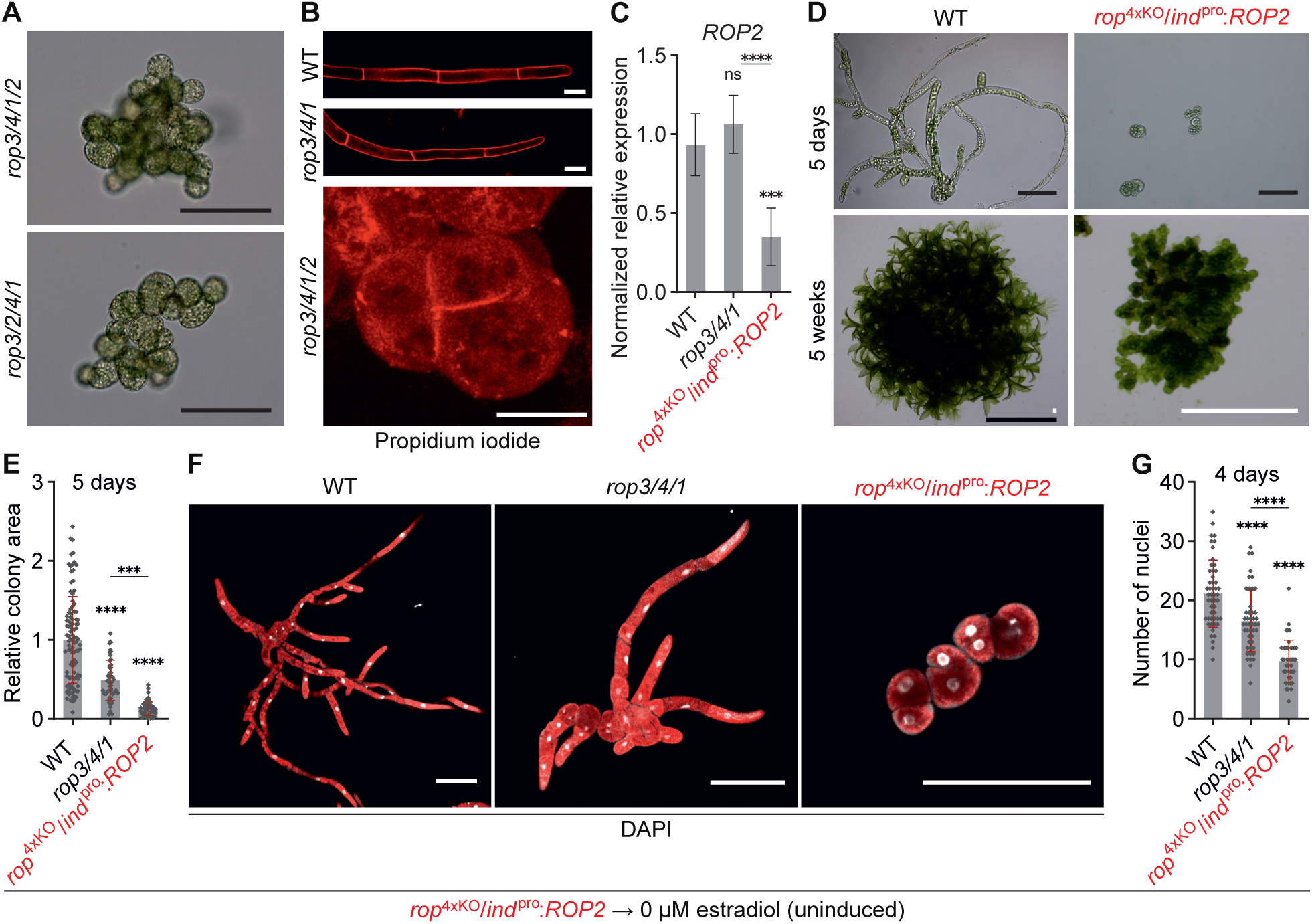
Loss of all Pp*ROP*s affects cell division and eliminates tip growth. Analysis of the *rop*^4xKO^ phenotype was performed using protonemata or 5-week-old colonies cultivated on media listed in Supplementary Table S4 with the indicated genotypes: i) WT, ii) *rop3/4/1,* iii) *rop3/4/1/2, rop3/2/4/1* (*rop*^4xKO^), iv) *rop*^4xKO^/*ind*^pro^:*ROP2* uninduced (conditional complementation of *rop*^4xKO^ with the coding and 3’ UTR sequence of Pp*ROP2* downstream of the inducible β-estradiol promoter). **A-C)** 1-week-old protonemata cultivated through homogenization. **A)** Bright field micrographs of *rop*^4xKO^ mutants. Scale bars: 100 µm. **B)** Confocal imaging of the cell division plane using projections of serial confocal optical sections of protonemata stained with propidium iodide. Scale bars: 25 µm. **C)** qRT-PCR of relative Pp*ROP2* transcript levels determined according to the 2^−ΔCCT^ method, using the value obtained for one WT replicate as a calibrator (relative expression = 1). Bars: mean of three biological and two technical replicates. The experiment was repeated two times with consistent results. **D-G)** Analysis of the *rop*^4xKO^ phenotype using 4 or 5-day-old protonemata or 5-week-old colonies regenerated from protoplasts with the indicated genotypes. *rop*^4xKO^/*ind*^pro^:*ROP2* protoplasts were regenerated without β-estradiol resembling the *rop*^4xKO^ phenotype. **D)** Bright field images from 5-day-old protonemata (top panel) and images of 5-week-old colonies (lower panel) were recorded using a stereo microscope. Scale bars: 100 µm (upper panel), 10 mm (lower panel, black bar), 500 µm (lower panel, white bars). **E)** Average size (area) of 5-day-old protonemata regenerated from protoplasts as determined based on microscopic imaging of chlorophyll autofluorescence using the mean value of WT as calibrator (relative area = 1). n = 50 colonies per genotype measured in 3 independent experiments. **F** and **G)** Cell division rate in 4-day-old protonemata was analyzed through projections of serial confocal optical sections displaying DAPI staining (white) and chlorophyll autofluorescence (red) (scale bars: 100 µm) **(F)** and determination of the average nuclei number **(G)**. n = 50 colonies per genotype measured in 3 independent experiments. **C, E, G)** Error bars: standard error of the mean (SEM) **(C)**, standard deviation (SD) **(E** and **G)**, dots represent individual data points. Statistical analysis by one-way ANOVA/Tukey’s test (pairwise comparisons to WT and relevant pairwise comparisons are displayed, all others see Supplementary Data Set S3): ^ns^ *P* > 0.05 (not significant), *** *P* ≤ 0.001, **** *P* ≤ 0.0001.

In addition to the defects described in the previous paragraph, quadruple Pp*ROP* RNAi knock-down mutants were demonstrated to also display reduced cell adhesion and altered cell wall composition (Burkart et al., 2015; Bao et al., 2022). Consistent with these observations, protoplasts could not be isolated from colonies formed by the two different *rop*^4xKO^ mutants described here, hampering the comparative quantitative phenotypic characterization of these mutants. To circumvent this issue, conditionally complemented *rop*^4xKO^ lines were generated. Based on homologous recombination the coding sequence (exons and introns) and 3′ UTR of the Pp*ROP3* gene in the *rop2/4/1* background was replaced by a CDS fragment encoding PpROP2, which was fused at 5’-end to an estradiol-inducible lexa/35S promoter (Kubo et al., 2013) and at the other end to the Pp*ROP2* 3′ UTR (Supplementary Figs. S2C: lower panel; and S3C). The two independent *rop*^4xKO^/*ind*^pro^:*ROP2* lines obtained based on this procedure were confirmed by PCR genotyping (Supplementary Fig. S3D). In the absence of estradiol, these lines produced Pp*ROP2* transcripts at substantially lower levels as compared to the *rop3/4/1* mutant (Fig. 4C), and displayed a typical *rop*^4xKO^ phenotype after 5 days and after 5 weeks in culture (Fig. 4D). Adding 10^-3^ µM estradiol to the culture medium resulted in partial complementation of the *rop*^4xKO^ phenotype, which enabled protoplast isolation.

Quantitative comparison of protoplast-derived 5-day-old protonemata growing on estradiol-free culture medium confirmed that *rop*^4xKO^/*ind*^pro^:*ROP2* colonies displaying a *rop*^4xKO^ phenotype were significantly smaller than *rop3/4/1* colonies (Fig. 4E), which showed the strongest growth defects of all *rop*^3xKO^ mutants when compared to WT (Fig. 2D and E). Remarkably, counting DAPI (4′,6-diamidino-2-phenylindole)-labeled nuclei in protoplast-derived protonemata grown for 4 days in the absence of estradiol (Fig. 4F) established that the observed stepwise reduction of the size of WT, *rop3/4/1* and *rop*^4xKO^/*ind*^pro^:*ROP2* (*rop*^4xKO^ phenotype) protonemata (Fig. 4E) is a consequence not only of inhibited cell expansion but also of a similar stepwise decrease in the rate of cell division (Fig. 4G). The cell division rate of apical protonemal cells is enhanced by caulonema differentiation (Cove and Knight, 1993; Jang and Dolan, 2011) and possibly by rapid cell expansion (Fantes and Nurse, 1977; Jones et al., 2017). The complete disruption of both these processes in *rop*^4xKO^ mutants (Fig. 4) and the partial inhibition of cell expansion in *rop*^3xKO^ mutants (Fig. 2), therefore appear likely to be partially responsible for the reduced cell proliferation displayed by these mutants. However, like related mammalian RHO GTPases (Coleman et al., 2004), PpROPs may also directly target cell cycle regulation to enhance the rate of cell division, in addition to promoting cell expansion and differentiation.

Comparing the phenotypes of *rop*^4xKO^ mutants and of the four *rop*^3xKO^ mutants expressing distinct individual PpROPs (Figs. 2-4) establishes that each of these proteins alone is capable of inducing not only normal polarization and directional expansion (albeit at the reduced rate) of protonemal cells, but also parallel cell division plane positioning, which is required for the development of linear filaments. Consistent with an important role of PpROP activity in cell division plane positioning, defects in this process were also observed a) in asymmetrically dividing cells in Arabidopsis roots overexpressing constitutively active mutant AtROP9 (Roszak et al., 2021) or in maize leaves lacking functional Zm*ROP2* and Zm*ROP9* genes (Humphries et al., 2011), b) during branch formation in *P. patens rop*^3xKO^ protonemata (Yi and Goshima, 2020), and c) in different tissues of the closely related liverwort *Marchantia polymorpha* after the disruption of the single Mp*ROP* gene expressed in this organism (Mulvey and Dolan, 2023b). Interestingly, quantitative phenotypic characterization of *rop*^3xKO^ and *rop*^4xKO^ mutants further demonstrated that PpROP activity directly and/or indirectly promotes the rate of cell division in growing protonemata. In summary, based on the investigation of these mutants PpROP activity was determined to control protonemal morphogenesis by promoting cell polarization, parallel cell division plane positioning, cell proliferation, and directional cell expansion.

### Estradiol-induced Pp*ROP2* expression at increasing levels progressively complements different defects displayed by *rop*^4xKO^ mutants

Data generated by the quantitative phenotypic characterization of *rop*^KO^ mutants (Figs. 1,2 and 4), and by the investigation of the ability of different PpROPs to complement *rop*^4xKO^ phenotypes (Fig. 3), strongly suggests that a) the four nearly identical PpROPs are functionally redundant, and b) increasing levels of total PpROP activity in apical protonemal cells are required for the normal polarization, division, expansion and differentiation of these cells (in this order). To further support these findings, the phenotypes of *rop*^4xKO^/*ind*^pro^:*ROP2* protonema were investigated, in which Pp*ROP2* expression was gradually induced by increasing estradiol concentrations in the culture medium. Estradiol concentrations eliciting striking phenotypic alterations were selected for this analysis (Fig. 5A). Control experiments confirmed that estradiol at these concentrations did not affect the development of WT protonemata (Supplementary Fig. S7).

**Figure 5.**
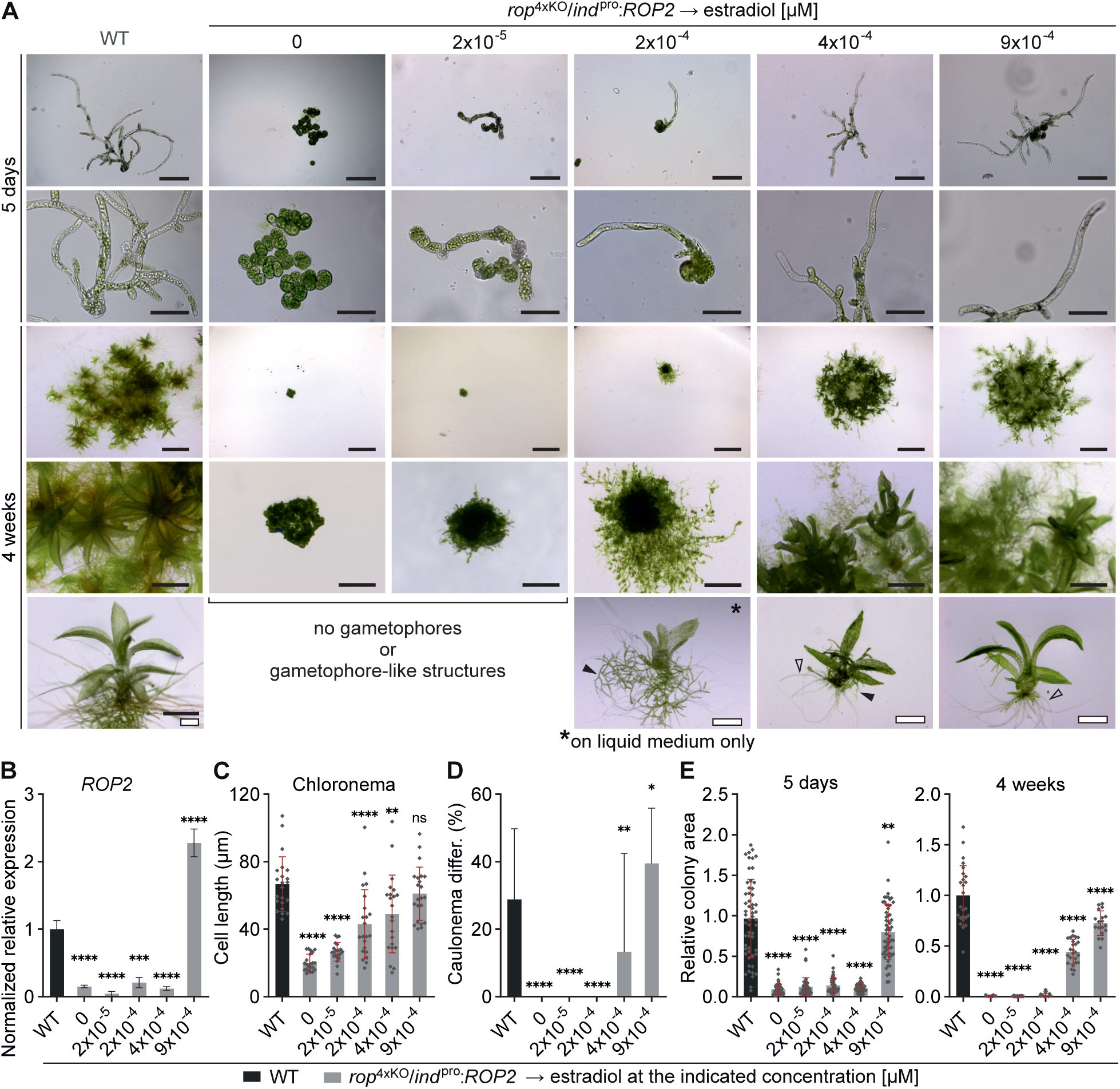
Titratable Pp*ROP2* expression rescues *rop*^4xKO^. Protonemata or 4-week-colonies were regenerated from protoplasts using media solidified with agar except where otherwise noted (see Supplementary Table S4) with the indicated genotypes: i) WT, ii) *rop*^4xKO^/*ind*^pro^:*ROP2* (conditional complementation of *rop*^4xKO^ with the coding and 3’ UTR sequence of Pp*ROP2* downstream of the inducible β-estradiol promoter) treated with representative β-estradiol concentrations. **A)** Bright field images of 5-day-old protonemata and images using a stereo microscope of 4-week-old colonies. *rop*^4xKO^/*ind*^pro^:*ROP2* did not develop gametophores (4 weeks) in the absence or with minimal PpROP2 expression (0 µM, 2x10^-5^ µM β-estradiol). Treatment with 2x10^-4^ µM β-estradiol generated gametophore-like structures only in liquid BCD medium (asterisk) and at ≥ 4x10^-4^ µM β-estradiol gametophores developed on BCD medium solidified with agar. Closed triangles indicate protonemata, open triangles indicate rhizoids. Scale bars: 200 µm (1^st^ top row), 100 µm (2^nd^ row), 5 mm (3^rd^ row), 1 mm (4^th^ row), 1 mm (5^th^ row, black bar), 500 µm (5^th^ row, white bars). **B)** Relative transcript levels of Pp*ROP2* were determined in 7-day-old protonemata (WT, *rop*^4xKO^/*ind*^pro^:*ROP2* treated with 9x10^-4^ µM β-estradiol) or 10-day-old protonemata (*rop*^4xKO^/*ind*^pro^:*ROP2* treated with ≤ 4x10^-4^ µM β-estradiol) according to the 2^-ΔΔCt^ method, using the value obtained for one WT replicate as calibrator (relative expression = 1). Bars: mean of three biological and two technical replicates. The experiment was repeated two times with consistent results. **C** and **D)** Average subapical cell length of chloronemata **(C)** or average percentage of caulonema differentiation as determined by microscopic observation **(D)** in 5-day-old protonemata. n = 22 cells **(C)** or n = 30 colonies **(D)** per genotype measured in 3 independent experiments. **E)** Average size (area) of 5-day-old or 4-week-old colonies using the mean value of WT as calibrator (relative area = 1). n = 60 colonies per genotype measured in three independent experiments (5 days). n = 22 colonies per genotype, the experiment was performed three times with consistent results (4 weeks). **B-E)** Error bars: standard error of the mean (SEM) **(B)**, standard deviation (SD) **(C-E)**. Dots represent individual data points. Statistical analysis by one-way ANOVA/Tukey’s test (pairwise comparisons to WT are displayed, all others see Supplementary Data Set S3): ^ns^ *P* > 0.05 (not significant), * *P* ≤ 0.05, ** *P* ≤ 0.01, *** *P* ≤ 0.001, **** *P* ≤ 0.0001.

At concentrations of up to 4x10^-4^ µM, estradiol-induced Pp*ROP2* expression remained much lower as compared to the WT expression level of this gene (Fig. 5B), which also corresponds to the level of total Pp*ROP* gene expression in *rop3/4/1* and in other *rop*^3xKO^ mutants (Fig. 3A, Supplementary Fig. S1). A linear correlation between estradiol concentrations of up to 4x10^-4^ µM and resulting minimal levels of Pp*ROP2* expression, which were near the detection limit, could not be established based on RT-qPCR analysis (Fig. 5B). Pp*ROP2* expression induced by 9x10^-4^ µM estradiol was about 2.3x stronger than in WT protonemata (Fig. 5B) and thus reached a level comparable to total Pp*ROP* gene expression in *rop*^2xKO^ mutants (2-3x WT Pp*ROP2* expression level; Supplementary Fig. S1). Unfortunately, higher estradiol concentrations failed to further enhance Pp*ROP2* expression to levels comparable to total Pp*ROP* gene expression in normally developing *rop*^1xKO^ or WT protonemata (3-4x or 5x WT Pp*ROP2* expression level, respectively; Supplementary Fig. S1).

As already shown above (Fig. 4D), in the absence of estradiol *rop*^4xKO^/*ind*^pro^:*ROP2* protonema displayed a typical *rop*^4xKO^ phenotype after 5 days and after several weeks in culture (Fig. 5). Remarkably, induction of Pp*ROP2* expression at the lowest level by 2x10^-5^ µM estradiol partially restored cell polarization and parallel cell division plane positioning required for filament formation (Fig. 5A), without substantially promoting directional cell expansion (Fig. 5C and E, Supplementary Table S1) or supporting caulonema differentiation (Fig. 5D). These observations demonstrate that cell polarization and parallel cell division plane positioning can be uncoupled from directional cell expansion, and that these processes depend on different levels of total PpROP activity.

On solid culture medium containing 2x10^-4^ µM estradiol, at low frequency chloronemal filaments were formed, which elongated based on directional cell expansion (Fig. 5A, Supplementary Table S1). However, this process was not fully restored, as subapical cells in these filaments were much shorter than corresponding WT cells (Fig. 5C), and colony growth remained strongly restricted (Fig. 5A and E). Caulonema differentiation (Fig. 5A and D) or gametophore formation (Fig. 5A) was not observed under these conditions. Interestingly, colonies growing for 4 weeks in liquid medium containing estradiol at the same concentration, and displaying an otherwise indistinguishable phenotype, frequently developed small gametophore-like structures, which contained two to four rudimentary phyllids, but failed to form stems or rhizoids (Fig. 5A). This finding is consistent with previous reports demonstrating that liquid culture can promote gametophore formation (Reski and Abel, 1985). As similar gametophore-like structures were never formed by *rop*^4xKO^/*ind*^pro^:*ROP2* colonies growing in liquid medium containing estradiol at lower concentrations, PpROP activity at minimal levels appears to be able to induce the formation of gametophore-like structures even in the absence of caulonema differentiation. Furthermore, PpROP activity at these levels evidently is sufficient for the development of rudimentary phyllids.

Pp*ROP2* expression induced by 4x10^-4^ µM estradiol further promoted directional cell expansion (Fig. 5A and C; Supplementary Table S1) and partially rescued caulonema differentiation (Fig. 5A and D). This led to a substantial increase in the size of 4-week-old colonies (Fig. 5A and E) and was accompanied by the frequent development of regular gametophores with a reduced size and severely stunted rhizoids on solid culture medium (Fig. 5A). However, the length of subapical protonemal cells, the rate of caulonema differentiation, as well as the size of 5-days-old and 4-weeks-old colonies remained significantly decreased. Interestingly, essentially the same developmental defects are shown by *rop*^3xKO^ mutants (Fig. 2), although the phenotype of these mutants appears to be somewhat weaker, as they display caulonema differentiation at WT rates, which are significantly reduced only in comparison to *rop*^2xKO^ mutants.

Finally, *rop*^4xKO^/*ind*^pro^:*ROP2* protonemata growing in medium containing 9x10^-4^ µM estradiol essentially phenocopied *rop*^2xKO^ mutants (Figs. 1 and 5, Supplementary Table S1), which as discussed above exhibit similar levels of total Pp*ROP* gene expression. As compared to WT protonemata, *rop*^4xKO^/*ind*^pro^:*ROP2* protonemata expressing Pp*ROP2* at this level of induction displayed significantly enhanced caulonema differentiation (Fig. 5D), while subapical cell length in 5-day-old protonemata was not detectably affected (Fig. 5C), and colony size after 5 days or after 4 weeks in culture as well as gametophore growth were only weakly reduced (Figs. 5A and E).

Data presented in this section demonstrate that PpROP2, a single member of the PpROP family, can effectively complement different severe defects displayed by *rop*^4xKO^ mutants. Partially interdependent developmental processes disrupted in these mutants were progressively restored by increasing levels of estradiol-induced Pp*ROP2* expression, indicating that these processes require distinct levels of total PpROP activity, rather than specific individual ROPs. Listed in the order of their increasing requirement for total PpROP activity, the following processes were restored by estradiol-induced PpROP2 expression: a) cell polarization and parallel cell division plane positioning required for filament formation, b) directional cell expansion, and c) development of gametophore-like structures or small gametophores, and d) caulonema differentiation.

### Pp*ROP* overexpression inhibits caulonema differentiation and depolarizes directional cell expansion

To investigate effects of PpROP overexpression, based on homologous recombination a CDS fragment encoding PpROP1, which was fused at the 5’-end to the estradiol-inducible lexa/35S promoter and at the other end to the Pp*ROP1* 3′ UTR, was introduced into a neutral region (Schaefer et al., 1991) of the WT genome (Supplementary Fig. S2D). One of the resulting WT/*ind*^pro^:*ROP1* lines, which displayed massive estradiol-inducible Pp*ROP1* overexpression, was confirmed by PCR genotyping (Supplemental Fig. S2E) and selected for phenotypic characterization. In the absence of estradiol, protoplast-derived protonemata of this line expressed Pp*ROP1* at WT level (Fig. 6A) and displayed no developmental defects (Fig. 6B-H). Growing these protonemata on solid medium containing 10^-3^ µM estradiol resulted in Rp*ROP1* expression at a more than 150x higher level (Fig. 6A), which roughly corresponds to a 30x increase in total Pp*ROP* gene expression as compared to WT protonemata (total Pp*ROP* gene expression ca. 5x stronger than Pp*ROP1* expression; Supplementary Fig. S1). Control experiments showed that estradiol at this concentration did not affect Pp*ROP1* expression or developmental processes in WT protonemata (Supplementary Fig. S7).

**Figure 6.**
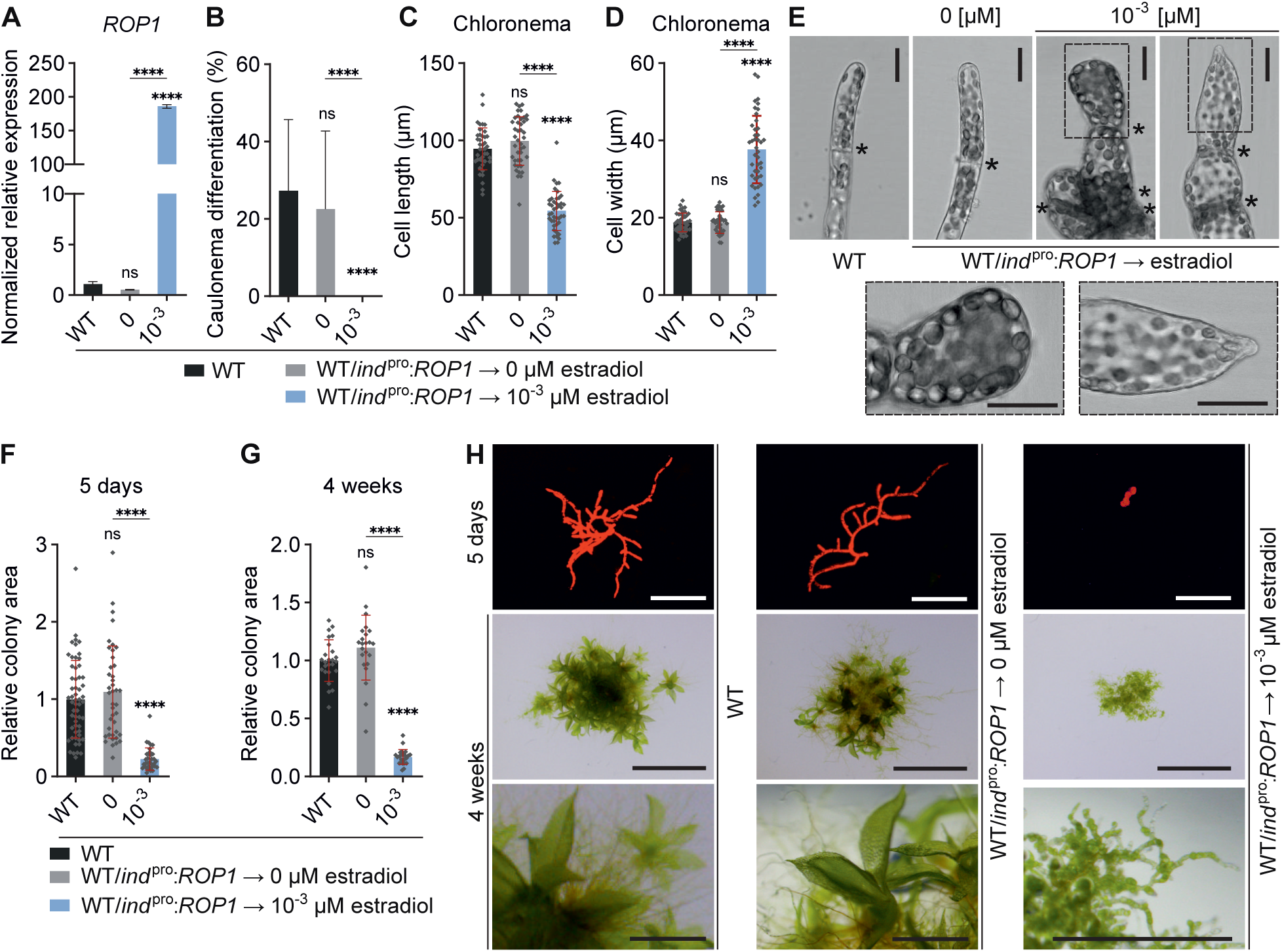
Pp*ROP* overexpression deregulates tip growth and caulonema differentiation. 1-week old protonemata cultivated through homogenization **(A)** or 5-day-old protonemata or 4-week colonies regenerated from protoplasts **(B-H)** using media listed in Supplementary Table S4 with the indicated genotypes: i) WT, ii) WT/i*nd^pro^*:*ROP1* (overexpression of the Pp*ROP1* coding sequence and 3’ UTR in WT using the inducible β-estradiol promoter. **A)** Relative transcript levels of Pp*ROP1* were determined according to the 2^−ΔCCT^ method using the value obtained for one WT replicate as calibrator (relative expression level = 1). Bars: mean of three biological and two technical replicates. The experiment was repeated two times with consistent results. **B-D)** Average percentage of caulonema differentiation as determined by microscopic observation **(B)** or average subapical chloronemal cell length **(C)** or cell width **(D)** in filaments with at least three cells of 5-day-old protonemata. n = 42 cells per genotype measured in 3 independent experiments. **E)** Bright field micrographs of filament tips of 5-day-old protonemata with the indicated genotypes. Asterisks indicate the cell wall between neighboring cells, dotted rectangles indicate the magnified apical region. Scale bars: 25 µm. **F-H)** Average size (area) of 5-day-old protonemata **(F)** or 4-week-old colonies **(G)** using the mean value of WT as calibrator (relative area = 1) were determined based on microscopic imaging of chlorophyll autofluorescence **(H**, 1^st^ top row; 5 days) or pictures recorded using a stereo microscope **(H**, 2^nd^ and 3^rd^ rows; 4 weeks). Scale bars: 500 µm (1^st^ top row), 5 mm (2^nd^ row), 1 mm (3^rd^ row). n = 35 **(F)** or n = 23 **(G)** colonies per genotype. The experiment was repeated two times, and the results from one representative experiment are shown here. **A-D, F, G)** Error bars: standard error of the mean (SEM) **(A)**, standard deviation (SD) **(B-D, F, G)**. Dots represent individual data points. Statistical analysis by one-way ANOVA/Tukey’s test (pairwise comparisons are displayed): ^ns^ *P* > 0.05 (not significant) **** *P* ≤ 0.0001.

As discussed above, PpROP activity was concluded to directly inhibit caulonema differentiation based on the observation that this process is enhanced in otherwise essentially normally developing *rop*^2xKO^ protonemata (Fig. 1). Further supporting this conclusion, Pp*ROP1* overexpression effectively disrupted caulonema differentiation (Fig. 6B). In angiosperm pollen tubes and root hairs excess ROP activity strongly depolarizes tip growth and results in massive apical ballooning (Chen et al., 2003; Gu et al., 2003a; Klahre and Kost, 2006; Kost, 2008). Overexpression of a fluorescent PpROP2 fusion proteins was reported to have similar but comparably weaker effects on directional expansion at the tip of *P. patens* protonemata (Ito et al., 2014). Quantitative analysis of protoplast-derived WT/*ind*^pro^:*ROP1* protonemata revealed that high-level Pp*ROP1* overexpression strongly reduced the length and enhanced the width of subapical cells in chloronemal filaments (Fig. 6C-E), which indicates massive depolarization of tip growth displayed by adjacent apical cells. Interestingly, PpPOP1 overexpression caused the apical dome of these cells to oscillate at irregular intervals between a wide flattened and a narrow pointed morphology (Fig. 6E). Together, inhibited caulonema differentiation and defective cell expansion strongly reduced the size of Pp*ROP1*-overexpressing 5-day-old protonemata and of 4-week-old colonies (Fig. 6F-H). Remarkably, WT/*ind*^pro^:*ROP1* protonemata never developed gametophores when growing on solid or in liquid medium containing 10^-3^ µM estradiol, demonstrating that PpPOP1 overexpression effectively blocked the formation of these structures (Fig. 6H).

In summary, consistent with and further defining PpROP functions identified by the phenotypic characterization of *rop*^KO^ mutants, PpROP1 overexpression prevented caulonema differentiation, effectively depolarized cell expansion at the tip of protonemal filaments, strongly reduced protonemal growth, and completely blocked gametophore formation. Overexpression of the PpROP effector PpRIC (ca. 40x WT Pp*RIC* expression level) also strongly inhibited caulonema differentiation and gametophore formation, but did not depolarize directional cell expansion and, consequently, only weakly reduced protonemal growth (Ntefidou et al., 2023). These observations strongly suggest that PpROP inhibits caulonema differentiation via PpRIC activation, but interacts with other effectors to control directional cell expansion. As discussed above, this conclusion is further supported by the increase or decrease in colony size displayed by *ric-1*^KO^ and *rop2x*^KO^ mutants, respectively. PpROP and PpRIC may also act together in a common pathway to directly inhibit gametophore formation, as this process is completely blocked by the overexpression of either of these two proteins. However, chloronemal filaments were proposed to lack competence for gametophore formation during normal protonemal development (Cove and Knight, 1993; Schumaker and Dietrich, 1997; Brun et al., 2003; Harrison et al., 2009), indicating that PpROP or PpRIC overexpression may indirectly block gametophore formation by inhibiting caulonema differentiation. Consistent with this interpretation, increasing ROP activity progressively stimulates the formation of gametophore-like structures and the development of gametophores (Figs. 2 and 5), suggesting that ROP activity promotes rather than inhibits these processes.

### PpROP functions in fundamental developmental processes depend on GDP/GTP cycling

Highly conserved amino acid residues can be mutated to alter the intrinsic rate of GDP/ GTP cycling displayed by RHO GTPases (Aspenström, 2019). These proteins are locked in the GTP-bound state by mutations corresponding to the Q61L exchange in H-RAS, which disrupt GTPase activity (GTP-locked). By contrast, mutations equivalent to the F28L exchange in H-RAS reduce nucleotide-binding affinity and enhance intrinsic GDP/GTP exchange (fast-cycling). Fast-cycling RHO mutants are preferentially GTP-bound at high GTP/GDP concentration ratios typically observed in the cytoplasm of eukaryotic cells. Like GTP-locked mutants, fast-cycling mutants are therefore considered to be constitutively active. To assess the importance of GDP/GTP cycling for PpROP functions in protonemal development, based on homologous recombination the coding sequence (exons and introns) and 3′ UTR of the endogenous Pp*ROP1* gene in the *rop3/2/4* background was replaced by CDS fragments encoding GTP-locked PpROP1^Q64L^ or fast-cycling PpROP1^F31L^, which were attached to the Pp*ROP1* 3′ UTR (Supplementary Fig. S2A and C: lower panel). Resulting *rop*^4xKO^/*ROP1*^pro^:*ROP1*^Q64L^ and *rop*^4xKO^/*ROP1*^pro^:*ROP1*^F31L^ lines were confirmed by PCR genotyping (Supplementary Fig. S2B). Based on RT-qPCR analysis of 1-week-old protonemata, lines expressing mutated Pp*ROP1* genes at the same level as Pp*ROP1* expression in WT plants and in *rop3/2/4* mutants were identified (Fig. 7A) and selected for phenotypic characterization (Fig. 7B-D).

**Figure 7.**
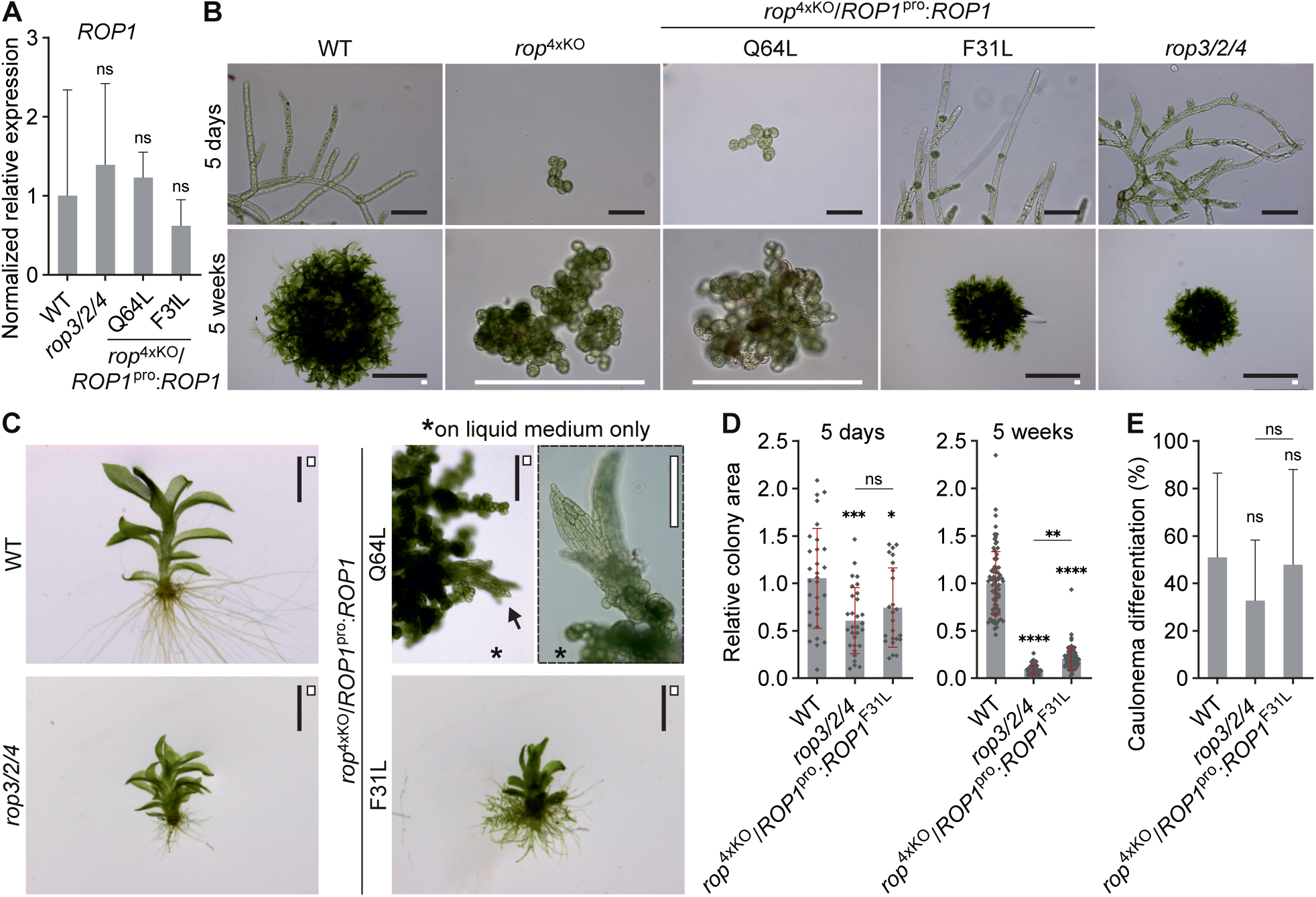
GTP/GDP cycling is essential for most PpROP functions but not gametophore initiation. Graphs and images based on protonemata or gametophores or colonies using culture conditions (Supplementary Table S4) with the indicated genotypes: i) WT, ii) *rop2/3/4*, iii) *rop*^4xKO^ iv) *rop*^4xKO^/*ROP1*^pro^:*ROP1*^Q64L^ (complementation of *rop*^4xKO^ with the Pp*ROP1* coding and 3’ UTR sequence containing a constitutive active mutation [Q64L] downstream of the endogenous Pp*ROP1* promoter), v) *rop*^4xKO^/*ROP1*^pro^:*ROP1*^F31L^ (complementation of *rop*^4xKO^ with the Pp*ROP1* coding and the 3’ UTR sequence containing a fast cycling mutation [F31L] downstream of the endogenous Pp*ROP1* promoter). **A)** Relative transcript levels of Pp*ROP1* in 1-week-old protonemata cultivated through homogenization according to the 2^−ΔCCT^ method, using the value obtained for one WT replicate as a calibrator (relative expression = 1). Bars: mean of three biological and two technical replicates. The experiment was repeated two times with consistent results. **B-E)** 5-day-old protonemata or 5-week-old colonies regenerated from protoplasts (WT, *rop2/3/4*, *rop*^4xKO^/*ROP1*^pro^:*ROP1*^F31L^) or cultivated through homogenization (*rop*^4xKO^, *rop*^4xKO^/*ROP1*^pro^:*ROP1*^Q64L^). **B)** Images of 5-day-old protonemata recorded using bright field microscopy (upper row) and images of 5-week-old colonies obtained using a stereo microscope (WT, *rop2/3/4*, *rop*^4xKO^/*ROP1*^pro^:*ROP1*^F31L^) or bright field microscopy (*rop*^4xKO^, *rop*^4xKO^/*ROP1*^pro^:*ROP1*^Q64L^) (lower row). Scale bars: 100 µm (upper row), 10 mm (black bars, lower row), 400 µm (white bars, lower row). **C)** Images of 5-week-old gametophores cultivated on solid BCD medium except *rop*^4xKO^/*ROP1*^pro^:*ROP1*^Q64L^ that developed gametophore-like structures only in liquid BCD medium (asterisk) were recorded using a stereo microscope. Arrow indicates a gametophore-like structure used for magnification using bright field microscopy (dotted rectangle). Scale bars: 2 mm (black bars) 200 µm (white bars). **D)** Average size (area) of 5-day-old protonemata or 5-week-old colonies normalized to the mean value of WT (relative area = 1). n = 30 colonies per genotype; the experiment was repeated three times with consistent results (5 days) or n = 81 colonies per genotype measured in 3 independent experiments (5 weeks). **E)** Average percentage of caulonema differentiation in 5-day-old protonemal filaments with at least three cells was determined by microscopic observation. n = 10 colonies per genotype. The experiment was repeated three times with consistent results. **A**, **D, E)** Error bars: standard error of the mean (SEM) **(A)**, standard deviation (SD) **(D** and **E),** dots represent individual data points. Statistical analysis by one-way ANOVA/Tukey’s test (pairwise comparisons to WT and relevant pairwise comparisons are displayed, all others see Supplementary Data Set S3). ^ns^ *P* > 0.05 (not significant), * *P* ≤ 0.05, ** *P* ≤ 0.01, *** *P* ≤ 0.001, **** *P* ≤ 0.0001.

Even after prolonged culture on solid medium, *rop*^4xKO^/*ROP1*^pro^:*ROP1*^Q64L^ colonies displayed a typical *rop^4x^*^KO^ phenotype (Fig. 7B). As this phenotype includes cell wall defects preventing protoplast isolation, *rop*^4xKO^/*ROP1*^pro^:*ROP1*^Q64L^ colonies shown were grown from tissues explants. This remarkable observation established that GDP/GTP cycling is strictly required for PpROP functions in fundamental developmental processes, including cell polarization and parallel division plane positioning required for filament formation. It remains to be determined whether GDP/GTP exchange is also essential for ROP functions in directional cell expansion and other processes, which appear to directly or indirectly depend on cell polarization and/or parallel division plane positioning. Interestingly, like *rop*^4xKO^/*ind*^pro^:*ROP2* colonies in the presence of estradiol at low concentration (2x10^-4^ µM; Fig. 5A), *rop*^4xKO^/*ROP1*^pro^:*ROP1*^Q64L^ colonies frequently formed gametophore-like structures exclusively when grown in liquid medium (Fig. 7C). The induction of gametophore formation in the absence of caulonema differentiation, as well as the promotion of rudimentary phyllid development by low-level PpROP activity, therefore does not require GDP/GTP cycling.

Protoplast-derived 5-day-old and 5-week-old *rop*^4xKO^/*ROP1*^pro^:*ROP1*^F31L^ and *rop3/2/4* protonemata displayed indistinguishable phenotypes (Fig. 7B). As compared to WT protonemata, colony size was similarly reduced (Fig. 7D), caulonema differentiation was not significantly affected (Fig. 7E), and gametophores with small but morphologically essentially normal leafy shoots and severely stunted rhizoids were formed (Fig. 7C). These data establish that by contrast to GTP-locked PpROP1^Q64L^, which is severely functionally impaired, fast-cycling PpROP1^F31L^ displays essentially the same ability as WT PpROP1 to promote protonemal development. However, it remains to be demonstrated that PpROP1^F31L^ is also capable of inhibiting caulonema differentiation when expressed at endogenous level in a *rop*^2xKO^ mutant, or upon overexpression.

### Overexpression of heterologous RHO GTPases in the *rop*^4xKO^ background suggests limited evolutionary conservation of PpROP downstream signaling

To test the ability of heterologous RHO GTPases to complement total loss of PpROP activity, AtROP7, the closest Arabidopsis PpROP homologue (Eklund et al., 2010), and human HsRHOA, one of the most extensively characterized RHO GTPases (Bishop and Hall, 2000), were overexpressed in the *rop*^4xKO^ background. To this end, based on homologous recombination, the coding sequence (exons and introns) of the endogenous Pp*ROP4* gene in the *rop3/1/2* background was replaced by CDS fragments encoding AtROP7 or HsRHOA, which were fused at 5’-end to the estradiol-inducible lexa/35S promoter (Supplementary Fig. S4A). Resulting *rop*^4xKO^/*ind*^pro^:At*ROP7* and *rop*^4xKO^/*ind*^pro^:Hs*RHOA* lines were confirmed by PCR genotyping (Supplementary Fig. S4B). Based on RT-qPCR analysis of 1-week-old protonemata (Fig. 8A), lines displaying maximal At*ROP7* or Hs*RHOA* transcript levels upon estradiol-induction were selected for phenotypic characterization (Fig. 8B and C). Maximal At*ROP7* and Hs*RHOA* transcript levels obtained were about 40x or more than 500x, respectively, higher than Pp*ROP4* transcript levels in WT plants and in *rop3/1/2* mutants (Fig. 8A), which roughly corresponds to 16x or more than 200x higher transcript levels as compared to total Pp*ROP* gene expression in WT protonemata (ca. 2,5x stronger than Pp*ROP4* expression; Supplementary Fig. S1). Control experiments showed that estradiol at concentrations of 10^-3^ µM or 1 µM, which maximally induced At*ROP7* or Hs*RHOA* expression, respectively, did not affect developmental processes in WT protonemata (Supplementary Fig. S7).

**Figure 8.**
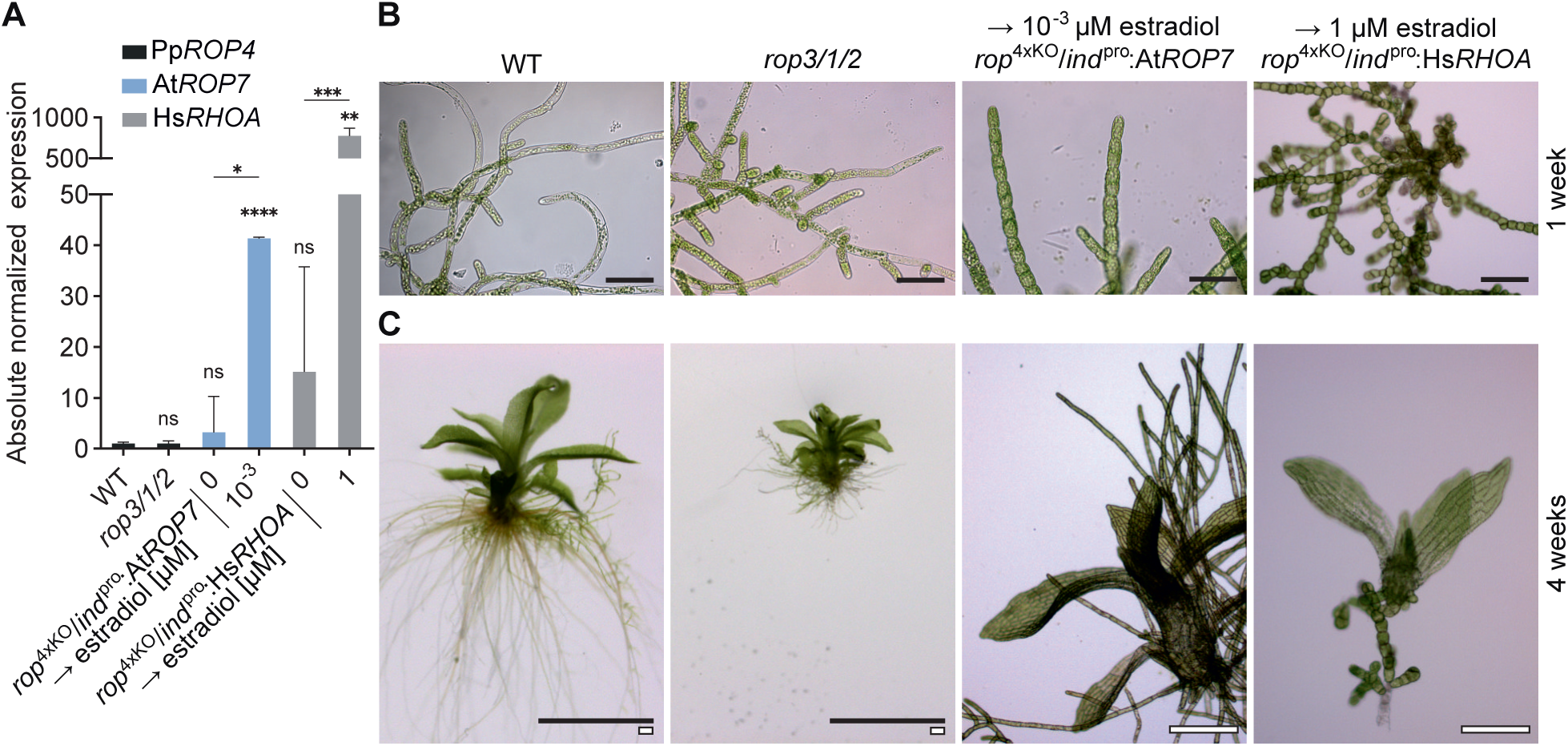
At*ROP7* or Hs*RHOA* overexpression partially rescues Pp*ROP*s. 1-week-old protonemata or 4-week-old colonies were cultivated through homogenization using media described in Supplementary Table S4 with the indicated genotypes: i) WT, ii) *rop1/2/3*, iii) *rop*^4xKO^/*ind*^pro^:At*ROP7* or *rop*^4xKO^/*ind*^pro^:Hs*RHOA* (complementation of *rop*^4xKO^ with the coding sequence of At*ROP7* or Hs*RHOA* downstream of the β-estradiol inducible promoter). **A)** Absolute transcript levels of Pp*ROP4*, At*ROP7,* or Hs*RHOA* were determined in 1-week-old protonemata with the indicated genotype based on standard curves, using the value obtained for one WT replicate of Pp*ROP4* as calibrator (relative expression = 1). Bars: mean of three biological and two technical replicates. Error bars: standard error of the mean. The experiment was repeated two times with consistent results. Statistical analysis by one-way ANOVA/Tukey’s test (pairwise comparisons to WT and relevant pairwise comparisons are displayed, all others see Supplementary Data Set S3): ^ns^ *P* > 0.05 (not significant), * *P* ≤ 0.05, ** *P* ≤ 0.01, *** *P* ≤ 0.001, **** *P* ≤ 0.0001. **B** and **C)** Bright field micrographs of 1-week-old protonemata **(B)**, or images of 4-week-old gametophores recorded using a stereo microscope (WT, *rop1/2/3*) or gametophore-like structures using a bright field microscope (*rop*^4xKO^/*ind*^pro^:At*ROP7*, *rop*^4xKO^/*ind*^pro^:Hs*RHOA*) **(C)**. Scale bars: 50 µm **(B)**, 2 mm (black bars), 200 µm (white bars) **(C)**.

Remarkably, massive At*ROP7* or Hs*RHOA* over-expression complemented the *rop*^4xKO^ phenotype to a similar extent (Fig. 8B and C) as estradiol-induced Pp*ROP2* expression at minimal level (2x10^-5^ µM; Fig. 5). Cell polarization and parallel cell division plane positioning required for filament formation were restored, whereas directional cell expansion was not substantially promoted and caulonema differentiation was never induced (Fig. 8B). Furthermore, even on solid culture medium, small gametophore-like structures occasionally developed, which contained a few rudimentary phyllids, but unlike regular WT or *rop*^3xKO^ gametophores failed to form stems or rhizoids (Fig. 8C). These observations establish that massive overexpression of heterologous flowering plant and mammalian RHO GTPases can restore fundamental developmental processes in *rop*^4xKO^ mutants, which require minimal levels of total PpROP activity. Consequently, PpROP downstream signaling controlling these processes appears to be evolutionary conserved at least to some extent. Supporting this conclusion, in *P. patens* KO mutants lacking geranylgeranyltransferase type I activity, which as discussed display a *rop*^4xKO^ phenotype, cell polarization and parallel cell division plane positioning could be restored by a YFP-AtROP1 fusion protein mutated to enable prenylation by farnesyltransferases. However, when tested using the same approach, human HsRAC1 or HsKRAS4b did not show a similar ability to complement loss of PpROP activity (Bao et al., 2022).

## Discussion

PpROP functions in protonemal development were systematically investigated based on quantitative phenotypic characterization a) of KO mutants lacking single Pp*ROP* genes or multiple such genes in all possible combinations, b) of complemented *rop*^4xKO^ lines expressing WT PpROPs, constitutively active PpROP1 mutants or heterologous RHO GTPases either at endogenous or different estradiol-titrated levels, and c) of lines displaying PpROP1 overexpression in the WT background. The results of these analyses firmly establish a) that the four nearly identical PpROPs, which are co-expressed at similar levels in protonemata, are functionally redundant, and b) that different developmental processes depend on distinct levels of total PpROP activity, rather than on specific individual ROPs (Fig. 9).

**Figure 9.**
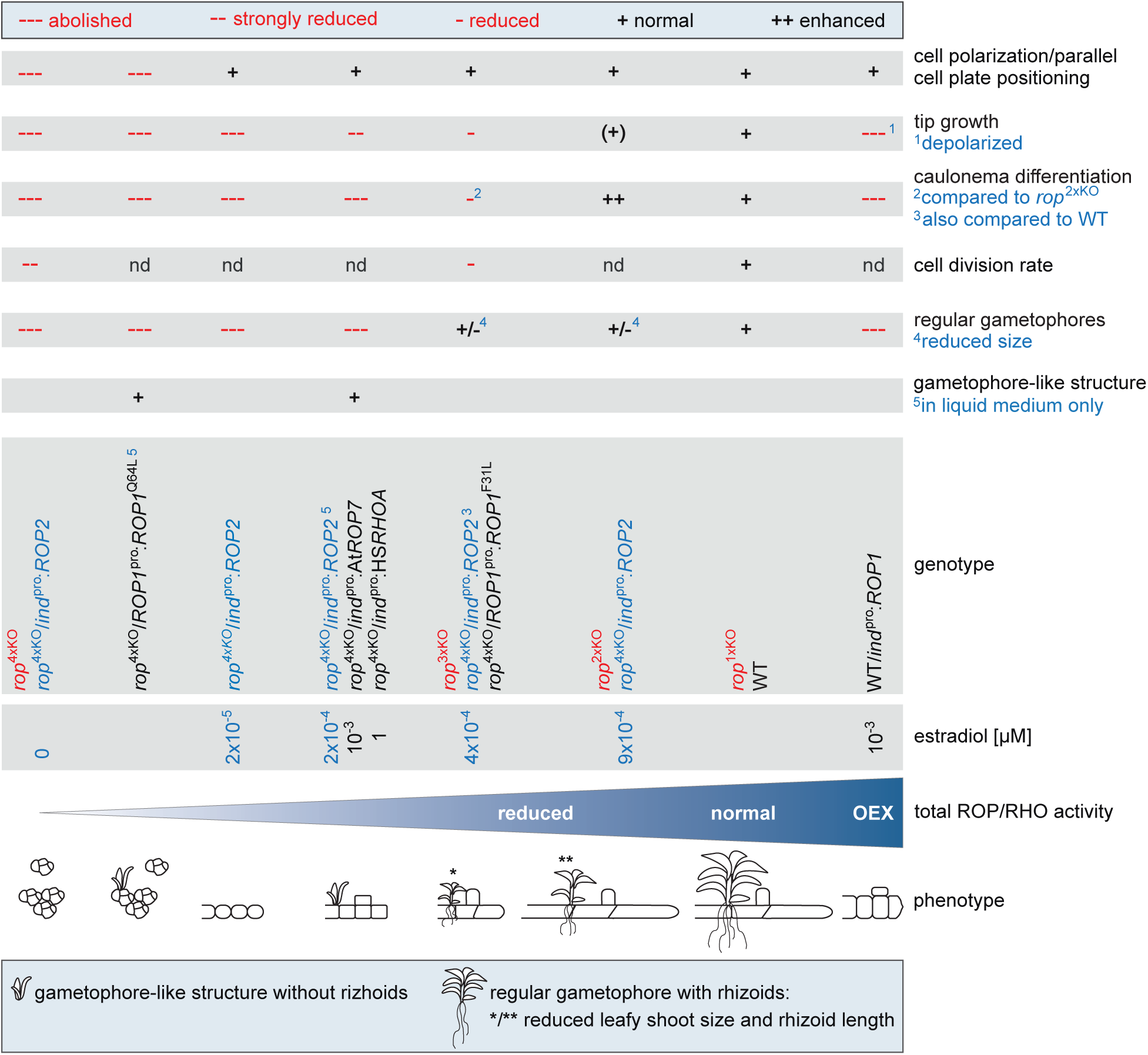
Overview of PpROP functions. Schematic drawing summarizing PpROP functions in protonemata and gametophores of *P. patens* with the indicated genotypes: i) WT (black font), ii) WT*/ind*^pro^:*ROP1* (Pp*ROP1* overexpresion in WT controlled by the inducible β-estradiol promoter) (Kubo et al., 2013) (black font), iii) *rop*^1xKO^*, rop*^2xKO^*, rop*^3xKO^, *rop*^4xKO^ (Pp*ROP* knockout mutants) (red font), iv) *rop*^4xKO^*/ROP1*^pro^:*ROP1*^Q64L^, *rop*^4xKO^*/ROP1*^pro^:*ROP1*^F31L^ (complementation of *rop*^4xKO^ with the coding and 3’ UTR sequence of Pp*ROP1* containing a constitutive active [Q64L] or fast cycling mutation [F31L] downstream of the Pp*ROP1* promoter) (black font), v) *rop*^4xKO^*/ind*^pro^:At*ROP7* or *rop*^4xKO^*/ind*^pro^:Hs*RHOA* (complementation of *rop*^4xKO^ with the coding sequence of At*ROP7* or Hs*RHOA* downstream of the inducible β-estradiol promoter) (black font), vi) *rop*^4xKO^*/ind*^pro^:*ROP2* (complementation of *rop*^4xKO^ with the coding and 3’ UTR sequence of Pp*ROP2* downstream of the inducible β-estradiol promoter) (blue font). nd: not determined.

### PpROP activity induces cell polarization and parallel division plain positioning at minimal levels, and increasingly promotes tip growth when further enhanced

Low levels of total PpROP activity induce cell polarization and parallel cell division plane positioning, which enables filament formation, but are insufficient for the promotion of directional cell expansion. Tip growth of apical protonemal cells is induced at somewhat higher total PpROP activity and is further promoted by increasing levels of this activity until it reaches WT rates in *rop*^1xKO^ mutants (Fig. 9). High-level PpROP overexpression massively depolarizes tip growth, similar to what has been observed in vegetative pollen tube cells and in root hairs (Chen et al., 2003; Gu et al., 2003a; Klahre and Kost, 2006; Kost, 2008). However, unlike these other cell types apical protonemal cells overexpressing PpROP activity do not simply swell at the tip, but display complex growth behavior that warrants further investigation (Fig. 6E). Despite these somewhat different overexpression effects, ROP functions in the control of tip growth appear to be essentially conserved from mosses to flowering plants. Yeast and mammalian RHO GTPases also have well-characterized functions in the regulation of directional cell expansion and related processes (Hall and Lalli, 2010; Ou and Yi, 2022).

### PpROP activity at WT and somewhat lower levels blocks caulonema differentiation via PpRIC stimulation

As caulonema differentiation is substantially enhanced both in *rop*^2xKO^ and *in ric-1*^KO^ mutants, total PpROP activity at levels reached in *rop*^1xKO^ and in WT protonemata appears to inhibit caulonema differentiation by stimulating the effector protein PpRIC (Fig. 9; Ntefidou et al., 2023). This conclusion is further supported by the recent demonstration that inhibition of PpROP activity by upstream regulators of the PpGAP (GTPase activating proteins) and PpREN (ROP Enhancer) families also promotes caulonema differentiation (Ruan et al., 2023). Consistent with these findings, caulonema differentiation is almost completely blocked by PpROP1 (Fig. 9) or PpRIC (Ntefidou et al., 2023) overexpression. However, as discussed above, apical chloronemal cells were proposed to undergo caulonema differentiation after reaching a threshold length (Jang and Dolan, 2011). Whereas PpRIC overexpression (ca. 40x WT Pp*RIC* expression level) does not substantially affect directional cell expansion (Ntefidou et al., 2023) and therefore appears to directly block caulonema differentiation, PpROP1 overexpression (ca. 30x increase in total Pp*ROP* gene expression as compared to WT; Fig. 6A) presumably interferes with this process also indirectly by disrupting tip growth. Similarly, the inhibition of caulonema differentiation observed at PpROP activity levels lower than those displayed by *rop*^2xKO^ mutants appears more likely to be a consequence of severely reduced directional cell expansion (Fig. 9) than to indicate a direct role of low-level PpROP activity in the stimulation of caulonema differentiation.

### PpROP activity contributes to the maintenance of apical initial cell identity

Protonemal filaments elongate based on tip growth and regular division of apical initial cells, which undergo gradual caulonema differentiation. PpROP activity appears to play an important role in the maintenance of apical initial cell identity not only by stimulating tip growth and by inhibiting caulonema differentiation as discussed above, but also by promoting the rate of cell division. Supporting this notion and consistent with previously reported functions of mammalian RHO GTPases in stimulating cell proliferation (Coleman et al., 2004), cell division rates were increasingly reduced in *rop*^3xKO^ and in *rop*^4xKO^ mutants (Fig. 9). As rapid cell expansion was proposed to enhance the proliferation of apical protonemal cells (Fantes and Nurse, 1977; Jones et al., 2017), PpROP activity may promote the division of these cells indirectly by increasing the rate of tip growth, rather than directly by accelerating cell cycle progression. Alternatively, PpROP activity may stimulate both these processes.

### PpROP activity induces the formation of gametophore-like structures and promotes the growth of regular gametophores

Regular gametophore formation was entirely abolished by high-level PpROP overexpression, as well as at levels of total PpROP activity lower than those displayed by *rop*^3xKO^ mutants (Fig. 9). The same conditions also completely blocked caulonema differentiation (Fig. 9), which was proposed to be essential for regular gametophore formation (Cove and Knight, 1993; Schumaker and Dietrich, 1997; Brun et al., 2003; Harrison et al., 2009). Strongly enhanced or reduced PpROP activity therefore appears to indirectly prevent regular gametophore formation by disrupting caulonema differentiation. However, in liquid culture medium estradiol-induced PpROP2 expression at low levels insufficient for regular gametophore formation, or PpROP1^Q64L^ expression at endogenous level, induced and supported the development of gametophore like-structures composed of rudimentary phyllids without rhizoids, which were never formed in the complete absence of PpROP activity (Fig. 9). Whereas regular gametophores develop from caulonema cells undergoing a well-defined series of asymmetric and symmetric cell divisions (Moody et al., 2018), gametophore like-structures emerged from undifferentiated or chloronemal cells dividing in a comparably random fashion (Fig. 9; Supplementary Fig. S8). Despite these differences, the observation that low-level PpROP activity can induce the formation of aberrant gametophore-like structures suggests that at higher levels enabling caulonema differentiation, this activity may also play an important role in initiating regular gametophore development. Furthermore, regular gametophore growth clearly depends on PpROP activity, since *rop*^2xKO^ *and rop*^3xKO^ mutants, as well as complemented *rop*^4xKO^ mutants displaying similar levels of total PpROP activity, formed gametophores with small leafy shoots and stunted rhizoids (Fig. 9). These observations demonstrate that PpROP activity not only has essential functions in promoting tip growth responsible for the elongation of protonemal filaments and rhizoids, but also plays an apparently somewhat less important role in stimulating other forms of directional cell expansion, which are required for the growth of leafy shoots.

### GDP/GTP cycling is essential for PpROP functions in fundamental developmental processes

When expressed at endogenous level in *rop*^4xKO^ mutants, both WT PpROP2 and fast-cycling PpROP1^F31L^ restored the *rop*^3xKO^ phenotype. By contrast, at the same expression level GTP-locked PpROP1^Q64L^ was unable to complement defects in cell polarization and parallel cell plate positioning displayed by *rop*^4xKO^ mutants, but induced the formation of gametophore-like structures by these mutants in liquid medium (Fig. 9). These observations demonstrate that GDP/GTP cycling is essential for PpROP functions in fundamental developmental processes underlying filament formation, but is not necessary for the induction of gametophore-like structures or directional cell expansion responsible for the growth of these structures. Further investigations are required to determine, whether GDP/GTP cycling is also essential for other PpROP functions such as the promotion of tip growth, which requires cell polarization, or the inhibition of caulonema differentiation, which depends on the formation of elongating filaments. While RHO GTPase functions generally appear to depend on GDP/GTP cycling, in animals atypical endogenous RHO GTPase have been identified, which like PpROP1^Q64L^ regulate selected cellular or developmental processes in a constitutively GTP-bound conformation (Hodge and Ridley, 2016; Aspenström, 2022).

### Evolutionary origin and maintenance of the Pp*ROP* gene family

*P. patens* protonemata express at similar levels four Pp*ROP* genes encoding nearly identical proteins, which as demonstrated by data presented here have essentially redundant functions and together provide total PpROP activity that governs key processes underlying protonemal development. A similar degree of sequence conservation and functional integration is not generally observed in eukaryotic gene families and raises questions concerning origin and maintenance of the Pp*ROP* gene family during evolution.

At least one *ROP* gene is generally expressed in all plants (viridiplantae) with the exception of some green algae of the chlorophyte group (Mulvey and Dolan, 2023b, a; Ntefidou et al., 2023). Interestingly, *ROP* gene families typically have between four and more than ten members in ferns, gymnosperm, and flowering plants, but are much smaller with only one or two members in most plants displaying more ancient features, which include lycophytes, the simplest vascular plants, along with mosses and all other non-vascular plants (Mulvey and Dolan, 2023a). Similarly, *RIC* gene families also substantially expanded and structurally differentiated with the emergence of flowering plants (Ntefidou et al., 2023). These observations suggest a) that the evolution of structurally increasingly complex vascular plants was accompanied by the diversification of ROP functions, and b) that one or two ROP isoforms are generally sufficient to exert comparably simple functions in plants with more ancient features. Consequently, total PpROP activity regulating *P. patens* protonematal development, which is provided by four co-expressed, nearly identical and functionally redundant PpROP family members, appears to correspond to the activity of one or two ROP isoforms in other non-vascular plants and in lycophytes.

The four members of the Pp*ROP* family are likely to have originated from a single ancient ROP gene as a consequence of two rounds of whole genome duplications (WGDs), which have occurred during the evolution of the *P. patens* genome (Rensing et al., 2007; Lang et al., 2018; Leebens-Mack et al., 2019; Gao et al., 2022). Supporting this notion, two consecutive WGDs and *ROP* gene families with four members have also been identified in mosses of the genus *Sphagnum*, whereas more than one WGD seems to have rarely occurred during the evolution of non-vascular plants with smaller *ROP* gene families (Supplementary Table S2; Lang et al., 2018; Leebens-Mack et al., 2019; Gao et al., 2022; Mulvey and Dolan, 2023b, a). The four *Sphagnum fallax* ROPs and all ROPs expressed in a number of mosses with smaller *ROP* gene families share at least 93 % identical amino acids with PpROP1 (Supplementary Table S2), suggesting that ROP functions are highly conserved in all mosses, which is in line with the hypothesis formulated at the end of the previous paragraph.

Remarkably, the four Pp*ROP* genes encoding nearly identical proteins, which apparently derived from genome duplications, were maintained in *P. patens* evidently in the absence of selective pressure driving their functional diversification. Although some degree of functional redundancy commonly appears to be evolutionary stable, particularly in gene families with important signaling functions (Vieten et al., 2005; Cevik et al., 2023; Kumar and Ruiz, 2023), the selective advantage of this phenomenon is not fully understood (Dean et al., 2008) and subject of ongoing further investigation (Nowak et al., 1997; Raval et al., 2023). Genomic functional redundancy can provide resilience against loss-of-function mutations (Gu et al., 2003b), as exemplified by the essentially normal development of *P. patens rop*^1xKO^ mutants. Furthermore, it increases gene dosage, which was proposed to enhance plant adaptability to changing environmental conditions (Kondrashov et al., 2002).

### Outlook

Evolutionary conserved essential functions of ROP GTPases in cell polarization, directional cell expansion and cell plate positioning have previously been established based on the investigation of defects caused by loss-of-function or gain-of-function mutations in the moss *P. patens* (Burkart et al., 2015; Cheng et al., 2020; Yi and Goshima, 2020; Bao et al., 2022), in the closely related non-vascular liverwort *M. polymorpha* (Mulvey and Dolan, 2023b), as well as in the flowering plants *A. thaliana* (Roszak et al., 2021) and *Z. mays* (Humphries et al., 2011). Data shown here demonstrate that during *P. patens* protonemal development PpROP activity has additional important functions in inhibiting caulonema differentiation, in controlling gametophore development, and in the direct or indirect regulation of cell proliferation. Whether ROP GTPases have similar or related functions also in other plants represents an important area of future research.

The discoveries summarized in the previous paragraph have provided an increasingly detailed understanding of ROP functions in the control of different cellular and developmental processes in plants. However, the signaling network stimulated by ROP activity to regulate these processes has not been well characterized. Compared to ROP and RHO GTPase functions, this network appears to be much less evolutionary conserved. In line with this notion, data presented here demonstrate that AtROP7 and HsRHOA can only complement the *rop*^4xKO^ phenotype to a very limited extent. Although these heterologous RHO GTPases are closely related to PpROPs (85.2-85.7 % [AtROP7] and 49.7 % [HsRHOA] amino acid identity) even upon massive overexpression they are only able to restore developmental processes in *rop*^4xKO^ mutants, which require minimal PpROP activity. Consistent with this finding, homologs of key animal RHO downstream effectors such as PAKs or ROCKs seem to be generally missing in plants. Instead, plant-specific ROP effectors including RICs, ICR/RIPs and RISAPs were shown to control directional cell expansion in flowering plants (Lavy et al., 2007; Yalovsky et al., 2008; Stephan et al., 2014). Interestingly, the *P. patens* genome encodes no ICR/RIP or RISAP homologs and only a single member of the RIC family (Eklund et al., 2010; Ntefidou et al., 2023). Furthermore, the single PpRIC protein is structurally very different from all its 11 *A. thaliana* homologs, and unlike several of these proteins does not regulate directional cell expansion, but inhibits caulonema differentiation (Ntefidou et al., 2023). Consequently, nothing is currently known about the effectors regulating tip growth downstream of PpROP activity in *P. patens*, and PpRIC is the only PpROP effector with a demonstrated function in the control of other cellular or developmental processes in this moss (Fig. 9). To further characterize the signaling network activated by PpROPs, it will be essential to identify and functionally characterize novel interaction partners and potential effectors of these proteins. The same approach will also provide important insights into the adaptation of ROP downstream signaling during plant evolution.

## Materials and Methods

### Plant materials and culture conditions

*P. patens* ecotype Gransden wild-type (WT) strain (2012) (Ashton and Cove, 1977) and transgenic lines (Supplementary Table S3) were grown axenically at 25°C under continuous white light illumination with fluorescent tubes (Philips Master TL-D Super 80 58 W, cool white) at an intensity of 50 µmol m^−2^ s^−1^. Unless otherwise stated, moss protonemata were cultivated and analyzed in 9-mm Petri dishes on BCDA medium (1 mM MgSO_4_, 1.84 mM KH_2_PO_4_, 10 mM KNO_3_, 5 mM ammonium-tartrate, 1 mM CaCl_2_, 45 µM FeSO_4_, 9.9 µM H_3_BO_3_, 2 µM MnCl_2_, 116 nM AlK[SO_4_]_2_, 424 nM CoCl_2_, 220 nM CuSO_4_, 235 nM KBr, 168 nM KI, 660 nM LiCl, 124 nM SnCl_2_, and 191 nM ZnSO_4_) (Ashton and Cove, 1977) solidified with 0.7% (w/v) agar and supplemented with 100 µg/mL vancomycin (Duchefa Biochemie) unless other antibiotics were used for transgene selection. Solidified medium was covered with cellophane discs (AA Packaging) (Grimsley et al., 1977) except for long-term cultivation of moss cultures consisting of small clumps of round cells. Protonemata were sub-cultured every 7 days and regenerated every 6 weeks from gametophores by homogenization (OMNI International) in Milli-Q H_2_O (Merck). Gametophore-forming colonies were cultivated and analyzed on BCD medium (BCDA without ammonium-tartrate, supplemented with 100 µg/ml vancomycin) solidified with 0.7% (w/v) agar or in liquid BCD medium. Titratable expression of RHO GTPases was induced by adding β-estradiol (Sigma), dissolved in DMSO at concentrations from 2x10^−5^ µM to 1 µM to BCDA or BCD medium after autoclaving. The culture conditions used in each experiment are listed in Supplementary Table S4.

### Phenotypic analysis of protonemata regenerated from protoplasts

Protonemata regenerated from protoplasts were prepared as described previously (Le Bail et al., 2019) based on Cove et al. (Cove et al., 2009). In brief, 4 to 7-day-old protonemata were digested with 0.5% (w/v) driselase (Sigma) in 8.5% mannitol and regenerated on PRMB medium (supplemented with 100 µg/ml vancomycin) solidified with 0.7 % agar in Petri dishes covered with cellophane for 2 days, followed by transfer of the cellophane disc to BCDA medium (supplemented with 100 µg/ml vancomycin). BCDA medium was treated with ≤ 1 µM β-estradiol for Pp*ROP1* overexpression (WT/*ind*^pro^:*ROP1*) or complementation of *rop*^4xKO^ with an inducible *ROP* or *RHO* expression construct (*rop*^4xKO^/*ind*^pro^:ROPX or *rop*^4xKO^/*ind*^pro^:*RHO*). Nuclei were counted 4 days after protoplast preparation. Cell length, caulonema differentiation and protonemal area were measured at 5 days. To analyze the gametophore stage, individual protonemal colonies, having regenerated for 5 days on BCDA were transferred to BCD medium without cellophane and supplemented with 100 µg/ml vancomycin, and the colony area was measured 4 or 5 weeks after protoplast preparation. The culture conditions used in each experiment are listed in Supplementary Table S4. Experiments were repeated three times. The cell length and the area of 5-day-old and 5-week-old colonies in Figures 1, 2, 4, and Supplementary Figure S5 were analyzed simultaneously.

### Bright field microscopy of cell length, caulonema differentiation, and colony size

To determine the cell length and caulonema differentiation, filaments with three or more cells of 5-day-old protonemata regenerated from protoplasts were imaged using a wide-field microscope (DMI4000B, Leica). The length of subapical cells was analyzed using Fiji software (ImageJ 1.54f; Schindelin et al., 2012), and caulonema differentiation was assessed by optical inspection of apical cells. The colony size was determined based on chlorophyll autofluorescence (excitation, 470 nm; emission, 500 nm long-pass) images obtained with a fluorescence stereo microscope (M205 FA, Leica) and analyzed using a Fiji macro (Vidali et al., 2007). 4 to 5-week-old colonies were imaged using the stereo microscope (M205 FA, Leica), and the area was determined by applying thresholding to convert reflected light 8-bit images to 1-bit images and analyzed using Fiji software (ImageJ 1.54f; Schindelin et al., 2012).

### Generation of knockout constructs

Molecular cloning was performed with standard methods (Green and Sambrook, 2012) using Phusion DNA polymerase (ThermoFisher Scientific), primers listed in Supplementary Data Set S1 (Eurofins Genomics), restriction enzymes and T4 Ligase (New England Biolabs). The resulting constructs (Supplementary Data Set S2) were verified by Sanger sequencing (Eurofins Genomics). To knock out Pp*ROP*s (*rop1*, *rop2*, *rop3*, and *rop4*) based on homologous recombination, genomic fragments of 0.7 to 1.0 kb upstream of the start codon (5′ target) and downstream of the stop codon (3′ target) were cloned flanking a resistance marker. The backbone of pMT123 (Thelander et al., 2004) was used for cloning into the XhoI/HindIII and SpeI/NotI sites, resulting in pSLU33 for knockout of Pp*ROP1* (Supplementary Fig. S2A and B); the p35S-Zeo backbone (Hiwatashi et al., 2008) was used for cloning pSLU34 for knockout of Pp*ROP2* (Supplementary Fig. S3A and B), the p35S-loxP-BSD backbone (Li et al., 2017) was used for cloning pSLU35 for knockout of Pp*ROP3* (Supplementary Fig. S3C and D), and the pMT164 backbone (Thelander et al., 2007) was used for cloning pSLU36 for knockout of Pp*ROP4* (Supplementary Fig. S4). Multiple Pp*rop* knockouts (*rop*^2xKO^, *rop*^3xKO^, *rop*^4xKO^) were generated through sequential knockout of single PpROPs in the order listed in Supplementary Fig. S2C. For each knockout, at least two lines were obtained except for *rop2*, *rop4*, and *rop4/1*, which had only one line each (Supplementary Table S3). All lines for each genotype showed the same phenotype, and the results of only one transgenic line are presented.

### Generation of expression constructs

To complement *rop*^4xKO^ with Pp*ROP1*^Q64L^ or Pp*ROP1*^F31L^ expressed by the endogenous Pp*ROP1* promoter, the Pp*ROP1* coding and 3’UTR sequence were amplified from cDNA and cloned into pENTR™/D-Topo (Thermo Fisher Scientific) yielding pFAU253 and mutated using PCR-based site-directed mutagenesis followed by DpnI digestion generating pFAU256 (Pp*ROP1*^Q64L^) or pFAU417 (Pp*ROP1*^F31L^). The Pp*ROP1*^Q64L^ or Pp*ROP1*^F31L^ fragments were transferred into the SalI/HindIII sites of pMT123 (Thelander et al., 2004) and flanked by 0.7 to 1.0 kb regions upstream of the start codon (5′ target) or downstream of the stop codon (3′ target) of Pp*ROP1* enabling the replacement of the Pp*ROP1* locus in *rop3/2/4* through homologous recombination yielding pFAU306 (*rop*^4xKO^/*ROP1*^pro^:*ROP1*^Q64L^) or pFAU421 (*rop*^4xKO^/*ROP1*^pro^:*ROP1*^F31L^) (Supplementary Fig. S2A and B; Supplementary Data Set S2).

To complement *rop*^4xKO^ with a single Pp*ROP* under the control of the Pp*ROP1* promoter, the full-length coding sequences of Pp*ROP2* or Pp*ROP3,* including the respective 3′ UTR, were amplified from cDNA and cloned into pENTR™/D-Topo, generating pFAU269 (Pp*ROP2*) and pFAU270 (Pp*ROP3*). Each Pp*ROP* fragment was then transferred from pFAU253 or pFAU269 or pFAU270 into the SalI/HindIII sites of pFAU306 which enabled the replacement of the Pp*ROP1* locus in *rop3/2/4* through homologous recombination, yielding pFAU508 (*rop*^4xKO^/*ROP1*^pro^:*ROP1*), pFAU509 (*rop*^4xKO^/*ROP1*^pro^:*ROP2*), and pFAU517 (*rop*^4xKO^/*ROP1*^pro^:*ROP3*) respectively (Supplementary Fig. S2A and B).

To complement *rop*^4xKO^ with a titratable expression of a *ROP* or *RHO GTPase*, the lexA operator and the XVE chimeric sequence were amplified by PCR from pGX8 (Kubo et al., 2013) and cloned into the NotI/EcoRV sites of pSLU35, yielding pFAU425, and into the SpeI/PmeI/PmlI sites of pSLU36, yielding pFAU391. The coding and 3’ UTR sequence of Pp*ROP2* were transferred from pFAU269 to pFAU425 by Gateway LR reaction (Thermo Fisher Scientific) (Katzen, 2007) generating pFAU431 (*rop*^4xKO^/*ind*^pro^:*ROP2*) (Supplementary Fig. S3C and D) that was used for replacement of the Pp*ROP3* locus based on homologous recombination in *rop2/4/1*. The full-length At*ROP7* and Hs*RHOA* coding sequences were amplified from Arabidopsis or *Homo sapiens* cDNA, respectively, and cloned into pENTR/D-Topo (Thermo Fisher Scientific), yielding pFAU290 (At*ROP7*) and pFAU294 (Hs*RHOA*), followed by Gateway LR reaction (Thermo Fisher Scientific) (Katzen, 2007) with pFAU391 generating pFAU398 (*rop*^4xKO^/*ind*^pro^:At*ROP7*) or pFAU402 (*rop*^4xKO^/*ind*^pro^:Hs*RHOA*) for gene replacement based on homologous recombination of the Pp*ROP4* locus in *rop3/1/2* (Supplementary Fig. S4A and B).

To overexpress Pp*ROP1* (WT/*ind*^pro^:*ROP1*) from the β-estradiol-inducible promoter, the coding sequence of Pp*ROP1*, including the 3′ UTR, was transferred from pFAU253 by Gateway LR reaction (Thermo Fisher Scientific) (Katzen, 2007) into pGX8 (Kubo et al., 2013) generating pFAU461 (Supplementary Fig. S2D) used for transgene integration based on homologous recombination into the neutral genomic region of PIG1b (Okano et al., 2009).

To express Pp*PINA*-*eGFP* under the endogenous Pp*PINA* promoter, the pFAUobt63 vector was obtained from Mattias Thelander (Viaene et al., 2014) for integration of *PINA*^pro^*:PINA*-*eGFP* into the neutral genomic region Pp108 (Schaefer and Zrÿd, 1997) of *P. patens* WT and *rop1/2* double knockout.

### *P. patens* transformation

Gene targeting into the *P. patens* genome by homologous recombination was previously described for polyethylene glycol PEG-mediated transformation (Le Bail et al., 2019) based on the method by Schaefer and Zrÿd (Schaefer and Zrÿd, 1997). In brief, 1.6 x 10^6^ protoplasts were transformed with 15 µg of linearized vector and allowed to regenerate for 5 days on PRMB medium, followed by two rounds of selection alternating between BCDA medium containing the appropriate antibiotic (30 µg/mL hygromycin B [Carl Roth], 20 µg/mL G418 [Merck], 50 µg/mL zeocin [Thermo Fisher Scientific], or 75 µg/mL blasticidin [InvivoGen]) according to the transformed resistance cassette and BCDA medium (supplemented with 100 µg/ml vancomycin). Transgenic lines were genotyped through PCR using primers indicated in Supplementary Data Set S1 and Supplementary Fig. S2-S4.

### Gene expression analysis by RT-qPCR

Total RNA was extracted from 1-week-old protonemata cultivated through homogenization or from 7 to 10-day-old protonemata regenerated from protoplasts. Samples were frozen in liquid nitrogen and lysed with glass beads using TissueLyser II (Qiagen). Total RNA was isolated using a Nucleospin RNA Plus^TM^ Kit (Macherey-Nagel), and RNA integrity was assessed using agarose gel electrophoresis. First-strand cDNA was synthesized from 1 µg of total RNA using iScript^TM^ Reverse Transcription Supermix for RT-qPCR (Bio-Rad) and diluted 1:20 in Milli-Q H_2_O. RT-qPCR was conducted using SscAdvanced^TM^ Universal SYBR Green Supermix (Bio-Rad) and a CFX96TM thermal cycler (Bio-Rad). Total RNA was prepared from three moss samples (biological replicates), and each cDNA sample was measured in two RT-qPCR reactions (technical replicates). Absolute transcript levels were quantified with standard curves prepared from a 10-fold dilution series of genomic DNA from 1-week old protonemata using the PhytoPure DNA Extraction kit (Cytiva). Pp*UBIQUITIN-E2* served as a reference gene (Le Bail et al., 2013). Primers used for RT-qPCR are listed in Supplementary Data Set S1. Experiments were repeated two times.

### Quantification of free IAA

7-day-old protonemata tissues (around 15-30 mg fresh weight per sample) were extracted after addition of 500 pg ^13^C6-IAA internal standard per sample, purified and analyzed for free IAA concentration using combined gas chromatography-tandem mass spectrometry as described (Andersen et al., 2008).

### Confocal microscopy and staining of nuclei and cell walls

A Leica TCS SP8 DIVE-FALCON inverted confocal laser scanning microscope with an HC PL APO CS2 20X/0.75 NA water immersion lens was used for confocal imaging, operated by Application Suite X software. GFP fluorescence was imaged through excitation using an argon laser at 488 nm and emission at 500-550 nm. The cell number was determined by counting nuclei in 4-day-old cells, or protonemata regenerated from protoplasts, stained according to the method of Vidali et al. (Vidali et al., 2010). In brief, moss samples were fixed for 30 min in a staining solution of 100 mM PIPES, pH 6.8, 0.1% (v/v) Nonidet P-40, 2% (w/v) paraformaldehyde, and 0.1 µg/mL 4′,6-diamidino-2phenylindole (DAPI). The fluorescence was detected through excitation using an argon laser at 360 nm and emission at 460 nm. Cell walls of gametophores were stained with 10 µg/mL propidium iodide solution (Merck), and the fluorescence was excited using a DPSS 561 laser (excitation, 536 nm; emission, 617 nm) and recorded using Z-stacks of optical sections at 2-µm spacing. Fiji software (Schindelin et al., 2012) was used to generate maximum Z-stack projections.

### Alignment analysis

A protein alignment of full-length PpROPs, AtROP7, and HsRHOA sequences and the protein identity matrix were obtained using Uniprot (The UniProt Consortium, 2024).

### Statistical analysis

GraphPad Prism (Version 10.3.1, GraphPad software) was used for data analysis. Unpaired Student’s *t*-tests were used to determine significant differences between two genotypes and one-way analysis of variance (ANOVA, Welch’s ANOVA) or two-way ANOVA followed by pairwise multiple comparison tests (Tukey’s/Dunnett’s) between three or more genotypes as indicated in the figure legends. Significance to WT and comparisons relevant to data interpretation are indicated in graphs, and all statistical results are given in Supplementary Data Set S3.

### Accession numbers

Sequence data were analyzed by BLAST (Altschul et al., 1990) and are available in the data libraries of Phytozome *P. patens* V6.1 (Joint Genome Institute) (Goodstein et al., 2012; Bi et al., 2024), TAIR (Arabidopsis Information Resource) (Berardini et al., 2015), ONEKP CNGBdb (Leebens-Mack et al., 2019), and GenBank (Clark et al., 2016) under the following accession numbers: Aa*ROP* (onekp|ZTHV_scaffold_2084304 Atrichum_angustatum, CNGBdb); Aat*ROP1* (onekp|QMWB_scaffold_2004844 Anomodon_attenuatus, CNGBdb); Aat*ROP2* (onekp|QMWB_scaffold_2004846 Anomodon_attenuatus, CNGBdb); Aat*ROP3* (onekp|QMWB_scaffold_2004845 Anomodon_attenuatus, CNGBdb); Ar*ROP* (gnl|onekp|WOGB_scaffold_2094631 Andreaea_rupestris, CNGBdb); At*ROP7* (At5g45970, TAIR); Ba*ROP* (gnl|onekp|JMXW_scaffold_2006533 Bryum_argenteum, CNGBdb); Bap*ROP* (gnl|onekp|HRWG_scaffold_2070290 Buxbaumia_aphylla, CNGBdb); Cc*ROP* (gnl|onekp|TAVP_scaffold_2005151 Calliergon_cordifolium, CNGBdb); Cd*ROP* (gnl|onekp|MIRS_scaffold_2008952 Climacium_dendroides, CNGBdb); Cp*ROP1* (gnl|onekp|FFPD_scaffold_2008289 Ceratodon_purpureus, CNGBdb); Cp*ROP2* (gnl|onekp|FFPD_scaffold_2009252 Ceratodon_purpureus, CNGBdb); Df*ROP* (gnl|onekp|AWOI_scaffold_2072508 Diphyscium_foliosum, CNGBdb); Fa*ROP* (gnl|onekp|DHWX_scaffold_2074728 Fontinalis_antipyretica, CNGBdb) Hs*RHOA* (NM_001664.4, GenBank); La*ROP* (gnl|onekp|VMXJ_scaffold_2010101 Leucobryum_albidum, CNGBdb); Ol*ROP1* (gnl|onekp|CMEQ_scaffold_2011516 Orthotrichum_lyellii, CNGBdb); Ol*ROP2* (gnl|onekp|CMEQ_scaffold_2013049 Orthotrichum_lyellii, CNGBdb); Pf*ROP* (gnl|onekp|ORKS_scaffold_2003761 Philonotis_fontana, CNGBdb); Pc*ROP* (onekp|SZYG_scaffold_2042265 Polytrichum_commune, CNGBdb); Pp*AUX1* (Pp6c16_1210, Phytozome); Pp*PINA* (Pp6c23_4450, Phytozome); Pp*ROP1* (Pp6c14_2140, Phytozome); Pp*ROP2* (Pp6c2_11900, Phytozome); Pp*ROP3* (Pp6c1_11060, Phytozome); Pp*ROP4* (Pp6c10_2660, Phytozome); Pp*RSL1 (*Pp6c1_20350, Phytozome); Pp*SHI1* (Pp6c21_9000, Phytozome); and Pp*UBIQUITIN-E2* (Pp6c12_2450, Phytozome); Rs*ROP1* (gnl|onekp|JADL_scaffold_2005567 Rhynchostegium_serrulatum, CNGBdb); Rs*ROP2 (gnl|onekp|JADL_scaffold_2005565 Rhynchostegium_serrulatum*, CNGBdb); Sf*ROP1* (Sphfalx06G097600, Phytozome); Sf*ROP2* (Sphfalx13G088000, Phytozome); Sf*ROP3* (Sphfalx14G076800, Phytozome); Sf*ROP4* (Sphfalx18G051700, Phytozome); Tl*ROP* (gnl|onekp|SKQD_scaffold_2079078 Takakia_lepidozioides, CNGBdb).

## Supporting information

Supplementary Data Sets S1-S3

## Author contributions

Designed the research: A.L.B, M.N., and B.K.; Performed research: A.L.B., M.N., J.N., T.I.L., D.K., K.L., H.V., S.S.; Data analysis: A.L.B., B.K., M.N., K.L.; Writing – original draft: M.N.; Writing – review & editing: B.K., M.N.; Visualization: M.N.; Supervision: B.K. and M.N.; Project administration: B.K.; Funding acquisition: K.L. and B.K. All authors are in agreement with this MS and approved its content.

## Acknowledgments

We are thankful to Dr. Mattias Thelander (Swedish University of Agricultural Sciences, Uppsala, Sweden) for the vector pMT123 and *PINA*^pro^:*PINA-GFP*, Prof. Mitsuyasu Hasebe (NIBB, Okazaki, Japan) for the vectors pGX8, p35S-Zeo, and p35S-loxP-BSD, and Dr. Pierre-François Perroud (IJPB, INRAE, Versailles) for invaluable technical and conceptual advice. We thank Roger Granbom for excellent technical support.

