## Supplementary Figures S1-S8, Tables S1-S4 for "Processes essential for *Physcomitrium patens* protonemal development require distinct levels of total activity provided by functionally redundant PpROP GTPases"

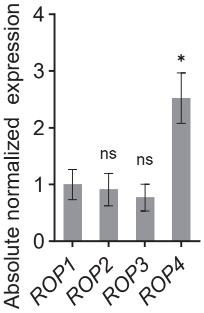


**Supplementary Figure S1. Pp*ROP* gene expression in protonemata** (supports **Figs. 1–5, 7**, **Supplementary Fig. S5**). Absolute transcripts levels of Pp*ROP*s in 1-week-old protonemata of WT cultivated through homogenization on BCDA medium (Supplementary Table S4) were determined based on standard curves, using the value obtained for one WT replicate of Pp*ROP1* as calibrator (relative expression = 1). Bars represent means of three biological and two technical replicates. Error bars: standard error of the mean (SEM). Statistical analysis by one-way ANOVA/Tukey's test (Supplementary Data Set S2): $^{ns}$ $P > 0.05$ (not significant); * $P \leq 0.05$.

**A**

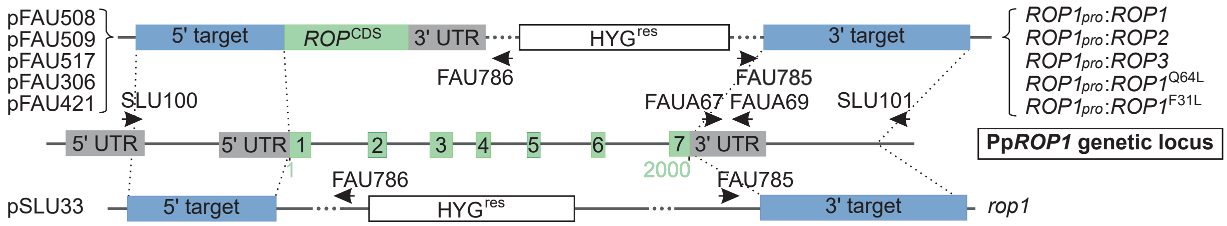


**B**

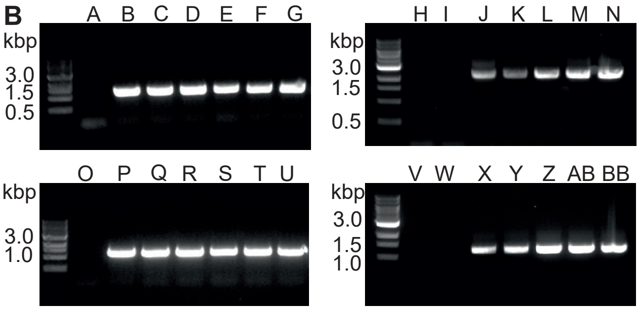


| lane | line | primers | bp |
|---|---|---|---|
| A | WT | FAU786+SLU100 | - |
| B | *rop1* | FAU786+SLU100 | 1279 |
| C | *rop3/1* | FAU786+SLU100 | 1279 |
| D | *rop4/1* | FAU786+SLU100 | 1279 |
| E | *rop2/4/1* | FAU786+SLU100 | 1279 |
| F | *rop3/4/1* | FAU786+SLU100 | 1279 |
| G | *rop3/2/4/1* | FAU786+SLU100 | 1279 |
| H | WT | FAU786+SLU100 | - |
| I | *rop3/2/4* | FAU786+SLU100 | - |
| J | *rop*$^{4xKO}$/*ROP1*$^{pro}$:*ROP1*$^{F31L}$ | FAU786+SLU100 | 2264 |
| K | *rop*$^{4xKO}$/*ROP1*$^{pro}$:*ROP1*$^{Q64L}$ | FAU786+SLU100 | 2264 |
| L | *rop*$^{4xKO}$/*ROP1*$^{pro}$:*ROP1* | FAU786+SLU100 | 2264 |
| M | *rop*$^{4xKO}$/*ROP1*$^{pro}$:*ROP2* | FAU786+SLU100 | 2264 |
| N | *rop*$^{4xKO}$/*ROP1*$^{pro}$:*ROP3* | FAU786+SLU100 | 2264 |
| O | WT | FAU785+SLU101 | - |
| P | *rop1* | FAU785+SLU101 | 1260 |
| Q | *rop3/1* | FAU785+SLU101 | 1260 |
| R | *rop4/1* | FAU785+SLU101 | 1260 |
| S | *rop2/4/1* | FAU785+SLU101 | 1260 |
| T | *rop3/4/1* | FAU785+SLU101 | 1260 |
| U | *rop3/2/4/1* | FAU785+SLU101 | 1260 |
| V | WT | FAU785+SLU101 | - |
| W | *rop3/2/4* | FAU785+SLU101 | - |
| X | *rop*$^{4xKO}$/*ROP1*$^{pro}$:*ROP1*$^{F31L}$ | FAU785+SLU101 | 1260 |
| Y | *rop*$^{4xKO}$/*ROP1*$^{pro}$:*ROP1*$^{Q64L}$ | FAU785+SLU101 | 1260 |
| Z | *rop*$^{4xKO}$/*ROP1*$^{pro}$:*ROP1* | FAU785+SLU101 | 1260 |
| AB | *rop*$^{4xKO}$/*ROP1*$^{pro}$:*ROP2* | FAU785+SLU101 | 1260 |
| BB | *rop*$^{4xKO}$/*ROP1*$^{pro}$:*ROP3* | FAU785+SLU101 | 1260 |

**C**

| *rop*$^{1xKO}$ | *rop*$^{2xKO}$ | *rop*$^{3xKO}$ | *rop*$^{4xKO}$ |
|---|---|---|---|
| *rop1* | *rop1/2* | | |
| *rop2* | *rop2/4* | *rop2/4/1* | |
| *rop3* | *rop3/1* | *rop3/1/2* | |
| | *rop3/2* | *rop3/2/4* | *rop3/2/4/1* |
| | *rop3/4* | *rop3/4/1* | *rop3/4/1/2* |
| *rop4* | *rop4/1* | | |

complemented *rop*$^{4xKO}$ lines

genetic background / transgene replacing remaining ROP gene

| *rop3/2/4* | *rop2/4/1* | *rop3/1/2* |
|---|---|---|
| *ROP1*$^{pro}$:*ROP1* | *ind*$^{pro}$:*ROP2* | *ind*$^{pro}$:AtROP7 |
| *ROP1*$^{pro}$:*ROP2* | | *ind*$^{pro}$:Hs*RHOA* |
| *ROP1*$^{pro}$:*ROP3* | | |
| *ROP1*$^{pro}$:*ROP1*$^{F31L}$ | | |
| *ROP1*$^{pro}$:*ROP1*$^{Q64L}$ | | |

**D**

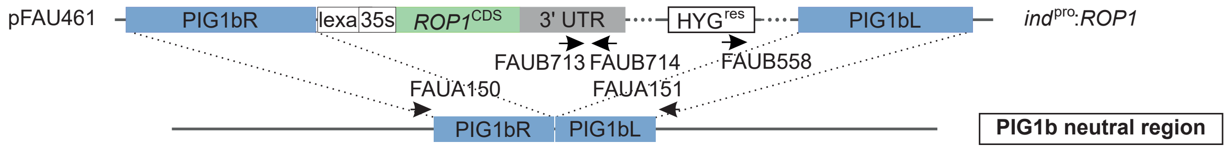


**E**

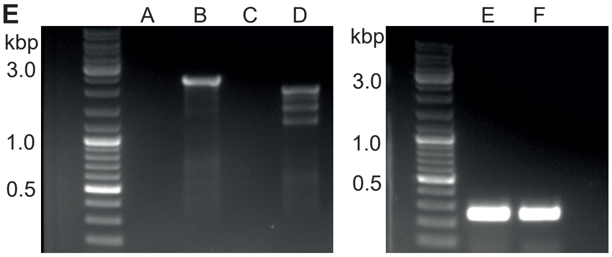


| lane | line | primers | bp |
|---|---|---|---|
| A | WT | FAUA150+FAUB714 | - |
| B | WT/*ind*$^{pro}$:*ROP1* | FAUA150+FAUB714 | 2429 |
| C | WT | FAUA151+FAUB558 | - |
| D | WT/*ind*$^{pro}$:*ROP1* | FAUA151+FAUB558 | 1987 |
| E | WT | FAUB713+FAUB714 | 251 |
| F | WT/*ind*$^{pro}$:*ROP1* | FAUB713+FAUB714 | 251 |

**Supplementary Figure S2. Editing of the genomic Editing of the genomic Pp*ROP1* locus through homologous recombination and summary of transgenic lines** (supports **Figs. 1–8, Supplementary Figs. S5, S6, S8**). **A)** Schematic representation of the Pp*ROP1* genetic locus and plasmid maps used to generate transgenic lines (Supplementary Table S3), drawn to scale. pSLU33 was used to generate *rop1* knockout by replacing the genomic fragment containing all exons and introns of Pp*ROP1* with the expression cassette, conferring resistance to hygromycin. pFAU508 (*ROP1*$^{pro}$:*ROP1*), pFAU509 (*ROP1*$^{pro}$:*ROP2*), pFAU517 (*ROP1*$^{pro}$:*ROP3*), pFAU306 (*ROP1*$^{pro}$:*ROP1*$^{Q64L}$) or pFAU421 (*ROP1*$^{pro}$:*ROP1*$^{F31L}$) were used to generate *rop*$^{4xKO}$ complementation lines by replacing all exons and introns of Pp*ROP1* in *rop3/2/4* with the coding and 3' UTR sequence of a Pp*ROP* variant. **A** and **D**) Green boxes: exons (numbered) or coding sequences, gray boxes: UTR sequences, blue boxes: regions used for homologous recombination targeting, white boxes: resistance markers, arrows: primers used for genotyping or RT-qPCR (Supplementary Data Set S1), three dots: sequence not displayed to scale to save space, green numbers: nucleotide position in the Pp*ROP1* coding region. **B)** Confirmation of transgenic lines through genotyping PCR using genomic DNA with the indicated primers. **C)** Summary of *rop* knockouts indicating the order in which they were generated from left to right (upper table). Summary of *rop*$^{4xKO}$ complementation lines generated by replacement of the remaining Pp*ROP* in the indicated *rop*$^{3xKO}$ genetic background (red font) with the coding sequence of an *ROP*/*RHO* gene expressed by the endogenous Pp*ROP1* promoter or the inducible β-estradiol-inducible promoter (bottom table). **D)** Vector map of pFAU461 used to overexpress *Pp**ROP1* in WT (WT/*ind*$^{pro}$:*ROP1*) by inserting the coding and 3' UTR sequence of Pp*ROP1* downstream of the inducible β-estradiol-inducible promoter in the PIG1 neutral region through homologous recombination. **E)** Confirmation of WT/*ind*$^{pro}$:*ROP1* transgenic line through genotyping PCR using genomic DNA with the indicated primers.

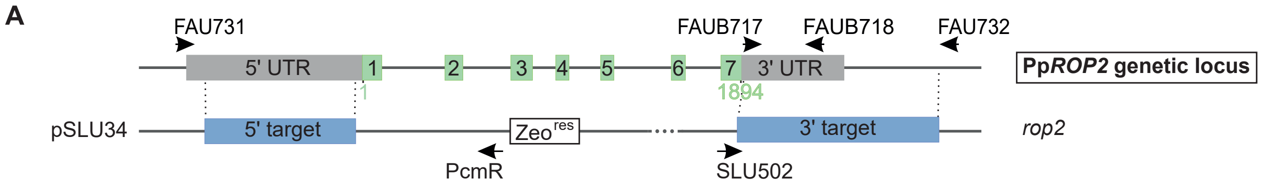


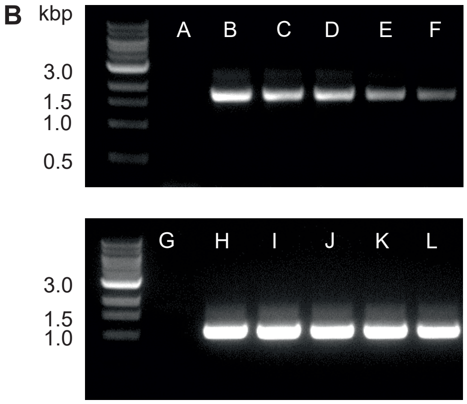


| lane | line | primers | size (bp) |
|---|---|---|---|
| A | WT | PcmR+FAU731 | - |
| B | *rop2* | PcmR+FAU731 | 1701 |
| C | *rop1/2* | PcmR+FAU731 | 1701 |
| D | *rop3/2* | PcmR+FAU731 | 1701 |
| E | *rop3/1/2* | PcmR+FAU731 | 1701 |
| F | *rop3/4/1/2* | PcmR+FAU731 | 1701 |
| G | WT | SLU502+FAU732 | - |
| H | *rop2* | SLU502+FAU732 | 1123 |
| I | *rop1/2* | SLU502+FAU732 | 1123 |
| J | *rop3/2* | SLU502+FAU732 | 1123 |
| K | *rop3/1/2* | SLU502+FAU732 | 1123 |
| L | *rop3/4/1/2* | SLU502+FAU732 | 1123 |

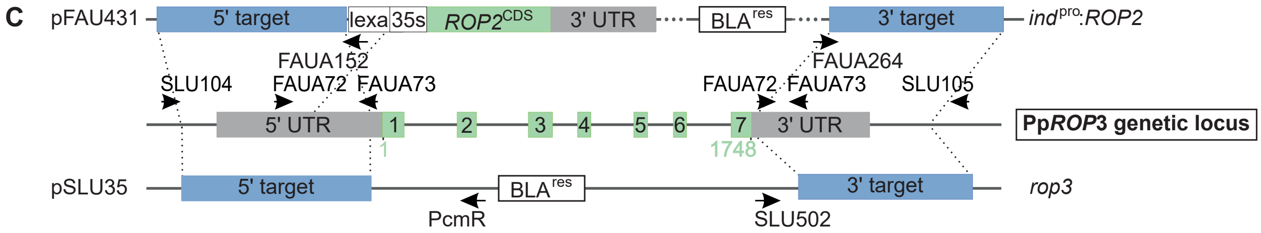


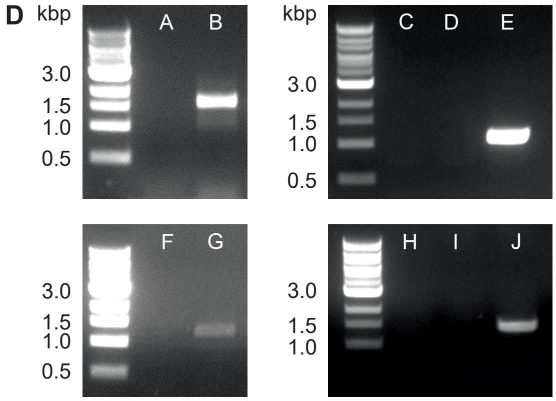


| lane | line | primers | size (bp) |
|---|---|---|---|
| A | WT | PcmR+SLU104 | - |
| B | *rop3* | PcmR+SLU104 | 1520 |
| C | WT | SLU104+FAUA152 | - |
| D | *rop2/4/1* | SLU104+FAUA152 | - |
| E | *rop*$^{4xKO}$/*ind*$^{pro}$:*ROP2* | SLU104+FAUA152 | 1083 |
| F | WT | SLU502+SLU105 | - |
| G | *rop3* | SLU502+SLU105 | 1251 |
| H | WT | SLU105+FAUA264 | - |
| I | *rop2/4/1* | SLU105+FAUA264 | - |
| J | *rop*$^{4xKO}$/*ind*$^{pro}$:*ROP2* | SLU105+FAUA264 | 1506 |

**Supplementary Figure S3. Editing of the genomic Pp*ROP2* and Pp*ROP3* loci through homologous recombination** (supports **Figs. 1–5**, **7, Supplementary Figs. S5, S6**). **A** and **C)** Schematic representation of the Pp*ROP2* **(A)** and Pp*ROP3* **(C)** genetic loci and plasmid maps used to generate transgenic lines (Supplementary Table S3), drawn to scale. Green boxes: exons (numbered) or coding sequences, gray boxes: UTR sequences, blue boxes: regions used for homologous recombination targeting, white boxes: resistance markers, arrows: primers used for genotyping or RT-qPCR (Supplementary Data Set S1), three dots: sequence not displayed to scale to save space, green numbers: nucleotide position in the Pp*ROP2* **(A)** or Pp*ROP3* **(C)** coding sequence. **A)** pSLU34 was used to generate *rop2* knockout by replacing the genomic fragment containing all Pp*ROP2* exons and introns with the expression cassette, conferring resistance to zeocin. **C)** pSLU35 was used to generate *rop3* knockout by replacing the genomic fragment containing all Pp*ROP3* exons and introns with the expression cassette, conferring resistance to blasticidin. pFAU431 was used to complement *rop*$^{4xKO}$ by replacing all exons and introns of the Pp*ROP3* locus with the coding and 3′ UTR sequence of Pp*ROP2* expressed under the control of the β-estradiol-inducible system generating *rop*$^{4xKO}$/*ind*$^{pro}$:*ROP2*. The lexA operator and 35S promoter are depicted, but not the GX8 promoter or XVE regions. **B** and **D)** Confirmation of transgenic lines through genotyping PCR using genomic DNA with the indicated primers.

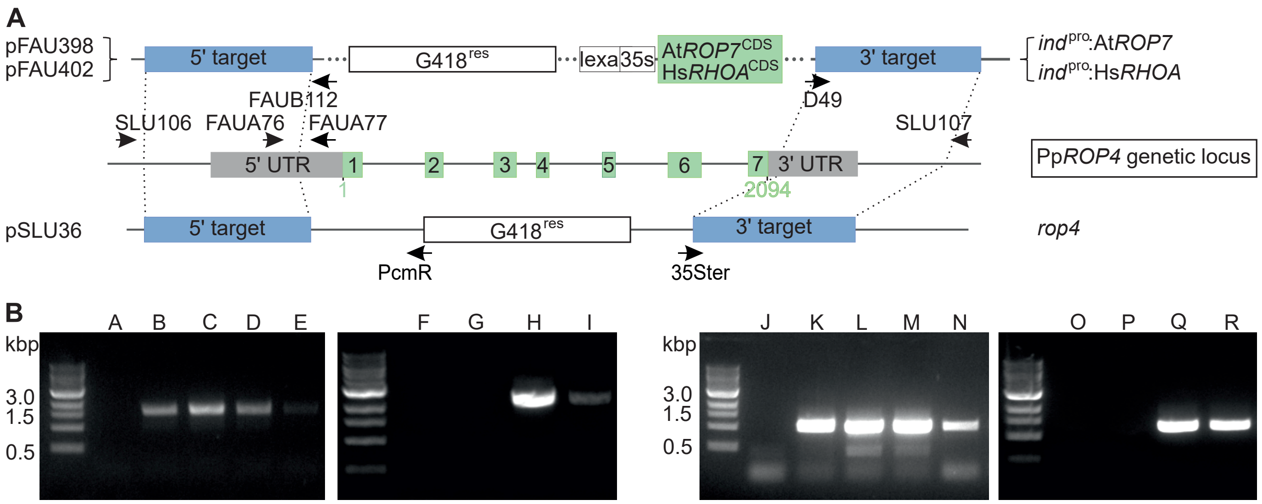


| lane | line | primers | size (bp) |
|---|---|---|---|
| A | WT | PcmR+SLU106 | - |
| B | *rop4* | PcmR+SLU106 | 1519 |
| C | *rop2/4* | PcmR+SLU106 | 1519 |
| D | *rop3/4* | PcmR+SLU106 | 1519 |
| E | *rop3/2/4* | PcmR+SLU106 | 1519 |
| F | WT | FAUB112+SLU106 | - |
| G | *rop3/1/2* | FAUB112+SLU106 | - |
| H | *rop*$^{4xKO}$/*ind*$^{pro}$:At*ROP7* | FAUB112+SLU106 | 2369 |
| I | *rop*$^{4xKO}$/ind$^{pro}$:Hs*RHOA* | FAUB112+SLU106 | 2369 |

| lane | line | primers | size (bp) |
|---|---|---|---|
| J | WT | 35Ster+SLU107 | - |
| K | *rop4* | 35Ster+SLU107 | 1018 |
| L | *rop2/4* | 35Ster+SLU107 | 1018 |
| M | *prop3/4* | 35Ster+SLU107 | 1018 |
| N | *rop3/2/4* | 35Ster+SLU107 | 1018 |
| O | WT | FAUD49+SLU107 | - |
| P | *rop3/1/2* | FAUD49+SLU107 | - |
| Q | *rop*$^{4xKO}$/*ind*$^{pro}$:At*ROP7* | FAUD49+SLU107 | 1260 |
| R | *rop*$^{4xKO}$/*ind*$^{pro}$:Hs*RHOA* | FAUD49+SLU107 | 1260 |

**Supplementary Figure S4. Editing of the genomic Pp*ROP4* locus through homologous recombination** (supports **Figs. 1–5**, **7**, **8, Supplementary Figs. S5, S6**). **A)** Schematic representation of the Pp*ROP4* genetic locus and plasmid maps used to generate transgenic lines (Supplementary Table S3), drawn to scale. pSLU36 was used to generate *rop4* knockout by replacing the genomic fragment containing all Pp*ROP4* exons and introns with the expression cassette, conferring resistance to G418. pFAU398 (*ind*$^{pro}$:At*ROP7*) or pFAU402 (*ind*$^{pro}$:Hs*RHOA*) were used to complement *rop*$^{4xKO}$ by replacing all exons and introns of Pp*ROP4* in *rop3/1/2* with the coding sequence of At*ROP7* or Hs*RHOA* expressed under the control of the β-estradiol-inducible system , generating *rop*$^{4xKO}$/*ind*$^{pro}$:At*ROP7* or *rop*$^{4xKO}$/*ind*$^{pro}$:Hs*RHOA*, respectively. Green boxes: exons (numbered) or coding sequences, gray boxes: UTR sequences, blue boxes: regions used for homologous recombination targeting, white boxes: resistance markers, arrows: primers (Supplementary Data Set S1) used for genotyping or RT-qPCR, three dots: sequence not displayed to scale to save space, green numbers: nucleotide position in the Pp*ROP4* coding sequence. **B)** Confirmation of transgenic lines through genotyping PCR using genomic DNA with the indicated primers.

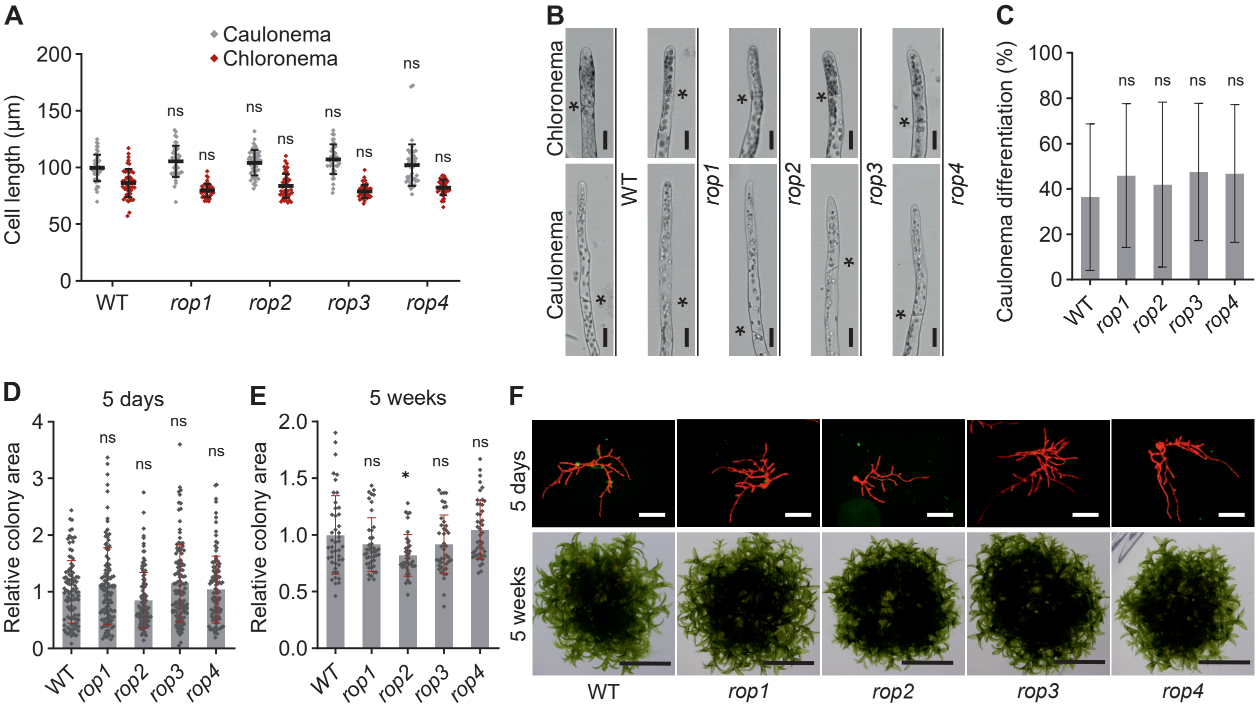


**Supplementary Figure S5**. **Knockout of a single Pp*ROP* (*rop*$^{1xKO}$) does not affect protonemata** (supports **Fig. 1**). **A–F)** Graphs and images based on 5-day-old protonemata or 5-week-old colonies regenerated from protoplasts were cultivated using media listed in Supplementary Table S4. **A)** Average subapical cell length of chloronemata and caulonemata in 5-day-old protonemata. n = 50 cells per genotype. The experiment was repeated three times with consistent results. **B)** Bright field micrographs of chloronemal and caulonemal filament tips. Asterisks: cell wall between apical and subapical cells. Scale bars: 25 µm. **C)** Average percentage of caulonema differentiation in 5-day-old protonema filaments with at least three cells as determined by microscopic observation. n = 60 colonies per genotype measured in 3 independent experiments. **D–F)** Average size of 5-day-old protonemata **(D)** or 5-week-old colonies **(E)** determined using micrographs of chlorophyll autofluorescence **(F**, upper row**)** or bright field images **(F**, lower row**)** recorded with a stereo microscope. n = 120 colonies per genotype measured in 3 independent experiments **(D)** or n = 41 colonies per genotype. The experiment was performed three times with consistent results **(E)**. Scale bars: 400 µm **(F,** upper row**),** 10 mm **(F,** lower row**)**. **A, C–E)** Error bars: standard deviation (SD), dots represent individual data points. Statistical analysis by two-way ANOVA/Tukey's test **(A)** or one-way ANOVA/Tukey's test **(C–E)**. Pairwise comparisons to WT are displayed, all others see Supplementary Data Set S2: $^{ns}$ *P* > 0.05 (not significant); * *P* ≤ 0.05.

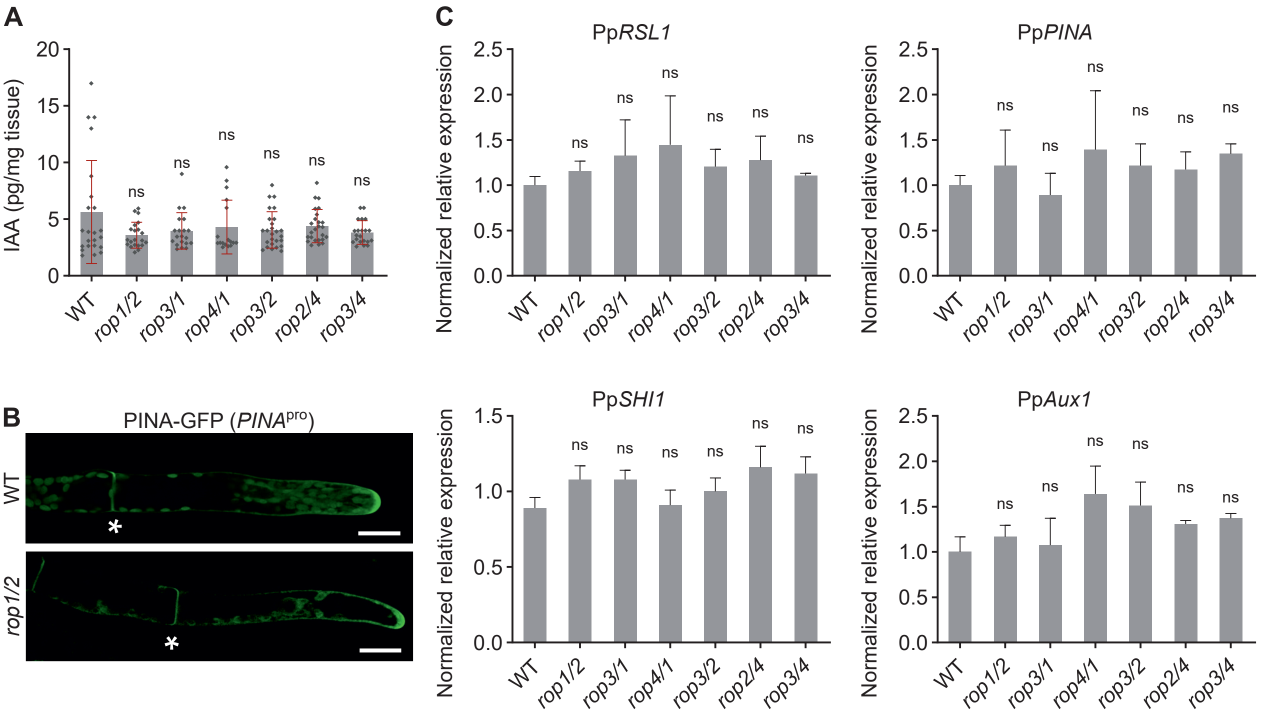


**Supplementary Figure S6**. **PpROPs do not influence the expression of auxin-regulated genes or the auxin content** (supports **Fig. 2**). **A–C)** Graphs and images based on 1-week-old protonemata with the indicated genotype cultivated using media described in Supplementary Table S4. **A)** Free IAA content of WT and *rop*$^{2xKO}$ lines were analyzed by gas chromatography - tandem mass spectrometry. Bars: means of n = 11 independent measurements, error bars: standard deviation, dots represent individual data points. **B)** Confocal microscopy imaging of PpPINA-GFP (Pp*PINA*$^{pro}$:Pp*PINA-GFP*) (Viaene et al., 2014) in WT and *rop1/2*. Asterisks indicate the cell wall between the apical and subapical cell. Scale bars: 25 µm. **C)** RT-qPCR was used to assess the expression of several auxin-responsive genes related to auxin signaling according to the $2^{-\Delta CCT}$ method, using the value obtained for one WT replicate as a calibrator (relative expression = 1). Bars: mean of three biological and two technical replicates, error bars: standard error of the mean (SEM). **A** and **C)** Statistical analysis by one-way ANOVA/Tukey's test (pairwise comparisons to WT are displayed, all others see Supplementary Data Set S2): $^{ns}$ $P > 0.05$ (not significant).

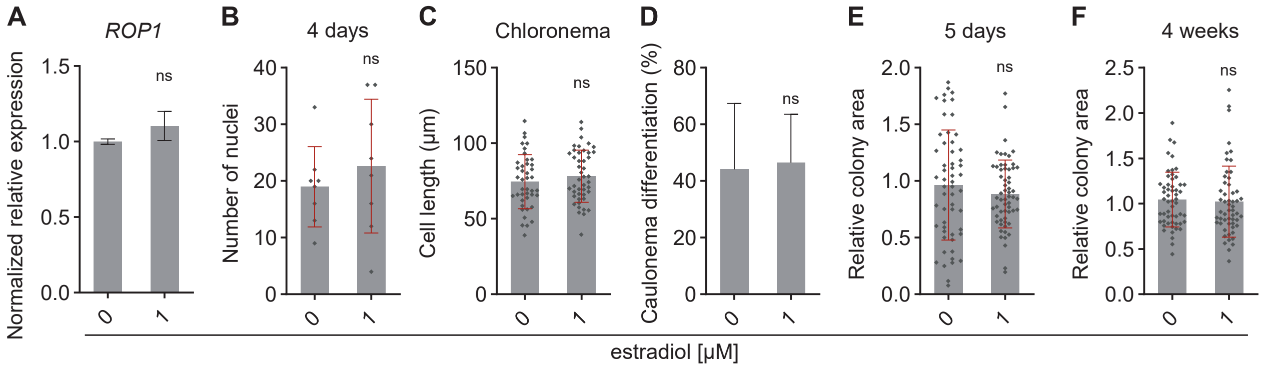


**Supplementary Figure S7. β-Estradiol does not influence Pp*ROP* expression or protonemal development** (supports **Figs. 5, 6, 8**). Graphs based on protonemata or 4-week-old colonies of WT cultivated using media listed in Supplementary Table S4. 1 µM β-estradiol was added to the media from a 10 mM stock solution dissolved in DMSO. **A)** Relative expression level of Pp*ROP1* in 1-week-old protonemata cultivated through homogenization was determined according to the $2^{-\Delta CCT}$ method, using the value obtained for one WT replicate as calibrator (relative expression = 1). Bars: mean of three biological and two technical replicates; error bars: standard error of the mean (SEM). The experiment was repeated two times with consistent results. **B–F)** Nuclei count and growth parameters were assessed using protoplasts regenerated for 2 days on PRMB medium without β-estradiol, followed by 2 days **(B)** or 3 days **(C–E)** on BCDA medium or 4 weeks **(F)** on BCD medium supplemented with 1 µM β-estradiol. **B)** Nuclei were counted in 4-day-old protonemata stained with DAPI using confocal fluorescence microscopy. n = 8 colonies per genotype. The experiment was repeated three times with consistent results. **C** and **D)** Average subapical cell length of chloronema cells **(C)** or average percentage of caulonema differentiation in 5-day-old protonemal filaments with at least three cells as determined by microscopic observation **(D)**. n = 42 cells per genotype were analyzed in 3 independent experiments **(C)**, or n = 20 colonies per genotype were analyzed. The experiment was repeated three times with consistent results **(D)**. **E** and **F)** Average size (area) of 5-day-old **(E)** or 4-week-old **(F)** colonies determined based on microscopic imaging of chlorophyll autofluorescence **(E)** or bright field micrographs **(F)** using a stereo microscope. The mean value of WT was used as a calibrator (relative area = 1). n = 55 protonemata **(F)** or n = 60 colonies **(E)** per genotype were analyzed in 3 independent experiments. **A–F**) Error bars: standard error of the mean (SEM) **(A),** standard deviation (SD) **(C–F)**; dots represent individual data points. Statistical analysis (Supplementary Data Set S2) by unpaired Student's *t*-test: $^{ns}$ $P > 0.05$ (not significant).

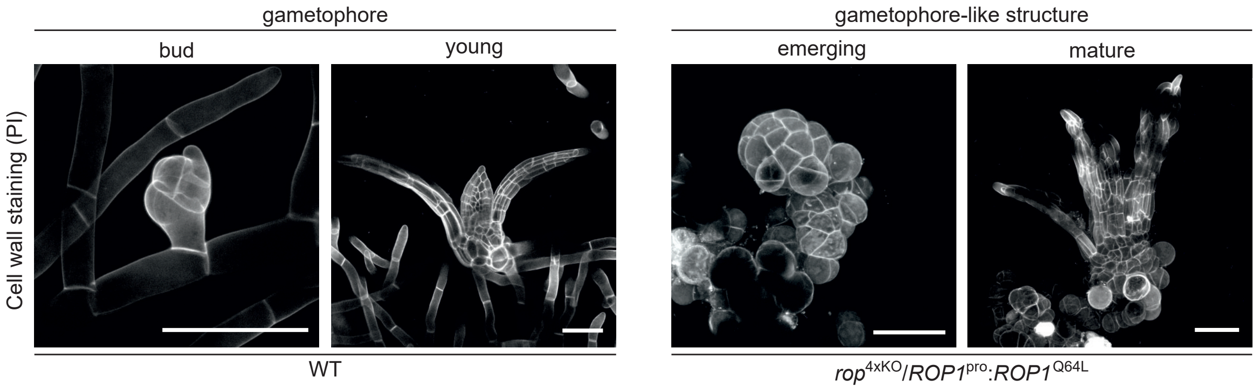


**Supplementary Figure S8. Gametophore-like structures of *rop*$^{4xKO}$/*ROP1*$^{pro}$:*ROP1*$^{Q64L}$** (supports **Fig. 7**). 5-week-old cultures were cultivated in liquid BCD medium (Supplementary Table S4). Maximum projections of serial optical sections of cell walls stained with propidium iodide were imaged using confocal microscopy. Scale bars: 100 µm.

**Supplementary Table S1**. **Rate of polarized cell expansion of *rop*$^{4xKO}$/*ind*$^{pro}$:*ROP2* at low induction level.** Cell elongation was assessed by optical inspection using a stereo microscope (M205 FA, Leica) in 5-day-old protonemata of *rop*$^{4xKO}$/*ind*$^{pro}$:*ROP2* regenerated from protoplasts on PRMB medium for 2 days followed by 3 days on BCDA medium supplemented with $2x10^{-5}$ to $9x10^{-4}$ µM β-estradiol. Detailed culture conditions listed in Supplemental Table S4. The experiment was repeated three times with consistent results.

| *rop*$^{4xKO}$/*ind*$^{pro}$:*ROP2* | | |
| --- | --- | --- |
| β-estradiol (µM) | 5-days-old protonemata with elongated cells (%) | n |
| 0 | 0 | 200 |
| $2x10^{-5}$ | 5 | 425 |
| $2x10^{-4}$ | 7 | 342 |
| $4x10^{-4}$ | 45 | 340 |
| $9x10^{-4}$ | 99 | 365 |

**Supplementary Table S2. ROP protein families in mosses: number of members and amino acid sequence conservation.** ROP protein families in mosses: number of members and amino acid sequence conservation. ROP homologs in selected moss species based on tBLASTN searches of Phytozome (Goodstein et al., 2012; Bi et al., 2024) and China National GeneBank Database (Leebens-Mack et al., 2019). Amino acid identity analyzed using tBLASTN (Altschul et al., 1990). Color code indicates moss species with 1 (white), 2 (blue), 3 (yellow), or 4 (green) ROP homologs. Whole genome duplication (WDG) data derived from [1]Rensing et al., 2007, [1]Lang et al., 2018, [2]Gao et al. 2022, [3]Leebens-Mack et al. 2019.

| **Species** | **ROP ID** | **Accession** | **Data library** | **Amino acid identity (%)** | | | | | | **WDG** |
|---|---|---|---|---|---|---|---|---|---|---|
| | | | | Pp_ ROP1 | Sf_ ROP1 | Cp_ ROP1 | Ol_ ROP1 | Ol_ ROP1 | Aa_ ROP1 | whole gemone duplication |
| *Physcomitrium patens* | PpROP1 | Pp6c14_2140 | Phytozome *P. patens* V6.1 | | | | | | | 2x WDG[1] |
| | PpROP2 | Pp6c2_11900 | | 99,5 | | | | | | |
| | PpROP3 | Pp6c1_11060 | | 99,5 | | | | | | |
| | PpROP4 | Pp6c10_2660 | | 100,0 | | | | | | |
| *Sphagnum fallax* | SfROP1 | Sphfalx06G097600 | Phytozome *S. fallax* V6.1 | 95,0 | | | | | | 2x WDG[2] |
| | SfROP2 | Sphfalx13G088000 | | 93,0 | 96,0 | | | | | |
| | SfROP3 | Sphfalx14G076800 | | 94,4 | 96,0 | | | | | |
| | SfROP4 | Sphfalx18G051700 | | 97,0 | 96,0 | | | | | |
| *Atrichum angustatum* | AaROP | onekp\|ZTHV_scaffold_2084304 Atrichum_angustatum | ONEKP CNGBdb | 99,0 | | | | | | 0 WDG[3] |
| *Andreaea rupestris* | ArROP | gnl\|onekp\|WOGB_scaffold_2094631 Andreaea_rupestris | ONEKP CNGBdb | 99,0 | | | | | | 0 WDG[3] |
| *Diphyscium foliosum* | DfROP | gnl\|onekp\|AWOI_scaffold_2072508 Diphyscium_foliosum | ONEKP CNGBdb | 99,0 | | | | | | 0 WDG[3] |
| *Takakia lepidozioides* | TlROP | gnl\|onekp\|SKQD_scaffold_2079078 Takakia_lepidozioides | ONEKP CNGBdb | 96,0 | | | | | | 0 WDG[3] |
| *Bryum argenteum* | BaROP | gnl\|onekp\|JMXW_scaffold_2006533 Bryum_argenteum | ONEKP CNGBdb | 99,0 | | | | | | 1x ancient WDG[3] |
| *Buxbaumia aphylla* | BapROP | gnl\|onekp\|HRWG_scaffold_2070290 Buxbaumia_aphylla | ONEKP CNGBdb | 100,0 | | | | | | 1x ancient WDG[3] |
| *Calliergon cordifolium* | CcROP | gnl\|onekp\|TAVP_scaffold_2005151 Calliergon_cordifolium | ONEKP CNGBdb | 99,0 | | | | | | 1x ancient WDG[3] |
| *Climacium dendroides* | CdROP | gnl\|onekp\|MIRS_scaffold_2008952 Climacium_dendroides | ONEKP CNGBdb | 99,0 | | | | | | 1x ancient WDG[3] |
| *Fontinalis antipyretica* | FaROP | gnl\|onekp\|DHWX_scaffold_2074728 Fontinalis_antipyretica | ONEKP CNGBdb | 99,0 | | | | | | 1x ancient WDG[3] |
| *Leucobryum albidum* | LaROP | gnl\|onekp\|VMXJ_scaffold_2010101 Leucobryum_albidum | ONEKP CNGBdb | 99,4 | | | | | | 1x ancient WDG[3] |
| *Philonotis fontana* | PfROP | gnl\|onekp\|ORKS_scaffold_2003761 Philonotis_fontana | ONEKP CNGBdb | 99,0 | | | | | | 1x ancient WDG[3] |
| *Polytrichum commune* | PcROP | onekp\|SZYG_scaffold_2042265 Polytrichum_commune | ONEKP CNGBdb | 98,0 | | | | | | 1x ancient WDG[3] |
| *Ceratodon purpureus* | CpROP1 | gnl\|onekp\|FFPD_scaffold_2008289 Ceratodon_purpureus | ONEKP CNGBdb | 99,0 | | | | | | 1x ancient WDG[3] |
| | CpROP2 | gnl\|onekp\|FFPD_scaffold_2009252 Ceratodon_purpureus | ONEKP CNGBdb | 99,0 | | 100,0 | | | | |
| *Orthotrichum lyellii* | OlROP1 | gnl\|onekp\|CMEQ_scaffold_2011516 Orthotrichum_lyellii | ONEKP CNGBdb | 98,0 | | | | | | 1x ancient WDG[3] |
| | OlROP2 | gnl\|onekp\|CMEQ_scaffold_2013049 Orthotrichum_lyellii | ONEKP CNGBdb | 97,0 | | | 97,0 | | | |
| *Rhynchostegium serrulatum* | RsROP1 | gnl\|onekp\|JADL_scaffold_2005567 Rhynchostegium_serrulatum | ONEKP CNGBdb | 97,0 | | | | | | 1x ancient WDG[3] |
| | RsROP2 | gnl\|onekp\|JADL_scaffold_2005565 Rhynchostegium_serrulatum | ONEKP CNGBdb | 97,0 | | | | 100,0 | | |
| *Anomodon attenuatus* | AatROP1 | onekp\|QMWB_scaffold_2004844 Anomodon_attenuatus | ONEKP CNGBdb | 99,5 | | | | | | 1x ancient WDG[3] |
| | AatROP2 | onekp\|QMWB_scaffold_2004846 Anomodon_attenuatus | ONEKP CNGBdb | 99,0 | | | | | 99,0 | |
| | AatROP3 | onekp\|QMWB_scaffold_2004845 Anomodon_attenuatus | ONEKP CNGBdb | 98,0 | | | | | 98,0 | |

**Supplementary Table S3**. ***P. patens* lines used in this study.** Moss lines and vectors used to generate transgene mutants through homologous recombination.

| ***P. patens* strain ID**<br>**(if applicable: β-estradiol levels used to induce transgene expression indicated in brackets: [ ])** | **Source** | **Vector ID** | **No. Of independent lines** |
|---|---|---|---|
| *Physcomitrium patens* ecotype Grandsen | Ashton and Cove, 1977 | N/A | - |
| *rop1*<br>(*rop1* knock-out mutant, homologous recombination) | this paper | pSLU33 | 3 |
| *rop2*<br>(*rop2* knock-out mutant, homologous recombination) | this paper | pSLU34 | 1 |
| *rop3*<br>(*rop3* knock-out mutant, homologous recombination) | this paper | pSLU35 | 2 |
| *rop4*<br>(*rop4* knock-out mutant, homologous recombination) | this paper | pSLU36 | 1 |
| *rop1/2*<br>(*rop1/2* knock-out mutant, homologous recombination) | this paper | pSLU33/pSLU34 | 5 |
| p*rop2/4*<br>(*rop2/4* knock-out mutant, homologous recombination) | this paper | pSLU34/pSLU36 | 3 |
| *rop3/1*<br>(*rop3/1* knock-out mutant, homologous recombination) | this paper | pSLU33/pSLU35 | 2 |
| *rop3/2*<br>(*rop3/2* knock-out mutant, homologous recombination) | this paper | pSLU34/pSLU35 | 5 |
| *rop3/4*<br>(*rop3/4* knock-out mutant, homologous recombination) | this paper | pSLU35/pSLU36 | 2 |
| *rop4/1*<br>(*rop4/1* knock-out mutant, homologous recombination) | this paper | pSLU33/pSLU36 | 1 |
| *rop2/4/1*<br>(*rop2/4/1* knock-out mutant, homologous recombination) | this paper | pSLU33/pSLU34/pSLU36 | 1 |
| *rop3/1/2*<br>(*rop3/1/2* knock-out mutant, homologous recombination) | this paper | pSLU33/pSLU34/pSLU35 | 3 |
| *rop3/2/4*<br>(*rop3/2/4* knock-out mutant, homologous recombination) | this paper | pSLU34/pSLU35/pSLU36 | 4 |
| *rop3/1/4*<br>(*rop3/1/4* knock-out mutant, homologous recombination) | this paper | pSLU33/pSLU35/pSLU36 | 2 |
| *rop3/2/4/1*<br>(*rop3/2/4/1* and *rop3/4/1/2* knock-out mutants, homologous recombination) | this paper | pSLU33/pSLU34/pSLU35/pSLU36 | 3 |
| *rop3/4/1/2*<br>(*rop3/2/4/1* and *rop3/4/1/2* knock-out mutants, homologous recombination) | this paper | pSLU33/pSLU34/pSLU35/pSLU36 | 1 |
| $rop^{4xKO}/ind^{pro}:ROP2_{cDNA}$<br>(*rop2/4/1/3* complemented with inducible Pp$ROP2_{cDNA}$ by gene replacement of Pp*ROP3* in *rop2/4/1*)<br>[β-estradiol: 0-1 μM] | this paper | pSLU33/pSLU34/pSLU36/pFAU431 | 2 |
| $rop^{4xKO}/ROP1^{pro}:ROP1_{cDNA}$<br>(*rop3/2/4/1* complemented with Pp$ROP1_{cDNA}$ by gene replacement of Pp*ROP1* in *rop3/2/4*) | this paper | pSLU34/pSLU35/pSLU36/pFAU508 | 3 |
| $rop^{4xKO}/ROP1^{pro}:ROP2_{cDNA}$<br>(*rop3/2/4/1* complemented with Pp$ROP2_{cDNA}$ by gene replacement of Pp*ROP1* in *rop3/2/4*) | this paper | pSLU34/pSLU35/pSLU36/pFAU509 | 3 |
| $rop^{4xKO}/ROP1^{pro}:ROP3_{cDNA}$<br>(*rop3/2/4/1* complemented with Pp$ROP3_{cDNA}$ by gene replacement of Pp*ROP1* in *rop3/2/4*) | this paper | pSLU34/pSLU35/pSLU36/pFAU517 | 2 |

| ***P. patens* strain ID**<br>**(if applicable: β-estradiol levels used to induce transgene expression indicated in brackets: [ ])** | **Source** | **Vector ID** | **No. Of independent lines** |
|---|---|---|---|
| *rop*$^{4xKO}$/*ROP1*$^{pro}$:*ROP1*$^{Q64L}$<br>(*rop3/2/4/1* complemented with Pp*ROP1*$^{Q64L}$ by gene replacement of Pp*ROP1* in *rop3/2/4*) | this paper | pSLU34/pSLU35/pSLU36/pFAU306 | 3 |
| *rop*$^{4xKO}$/*ROP1*$^{pro}$:*ROP1*$^{F31L}$<br>(*rop3/2/4/1* with Pp*ROP1*$^{F31L}$ by gene replacement of Pp*ROP1* in *rop3/2/4*) | this paper | pSLU34/pSLU35/pSLU36/pFAU421 | 4 |
| *rop*$^{4xKO}$/*ind*$^{pro}$:At*ROP7*$_{cDNA}$<br>(*rop3/1/2/4* complemented with inducible At*ROP7* by gene replacement of Pp*ROP4* in *rop3/1/2*)<br>[β-estradiol: 0-10$^{-3}$ µM] | this paper | pSLU33/pSLU34/pSLU35/pFAU398 | 2 |
| *rop*$^{4xKO}$/*ind*$^{pro}$:Hs*RHOA*$_{cDNA}$<br>(*rop3/1/2/4* complemented with inducible Hs*RhoA* by gene replacement of Pp*ROP4* in *rop3/1/2*)<br>[β-estradiol: 0-1 µM] | this paper | pSLU33/pSLU34/pSLU35/pFAU402 | 2 |
| WT/*ind*$^{pro}$:*ROP1*$_{cDNA}$<br>(inducible Pp*ROP1* targeted in PIG1 neutral region of WT)<br>[β-estradiol: 0-1 µM] | this paper | pFAU461 | 4 |
| *PINA*$^{pro}$:*PINA-GFP*<br>(homologous recombination in WT P108 neutral genomic locus) | this paper | pFAUobt63 | 2 |
| *rop*$^{2xKO}$/*PINA*$^{pro}$:*PINA-GFP*<br>(homologous recombination in P108 neutral genomic locus of *rop1/2*) | this paper | pSLU33/pSLU35/pFAUobt63 | 2 |

**Supplementary Table S4. Culture media used in this study.** *P. patens* was cultivated using BCD or BCDA solidified with agar or liquid media (Asthon and Cove, 1977) and supplemented with vancomycin (bactericidal antibiotic). β-estradiol was added to media for inducible gene expression using the XVE system (Kubo et al., 2013). Protonemata or gametophores used in experiments were obtained through homogenization or regeneration from protoplasts. Color coding: figures (black font), supplementary figures (red font), supplementary table (green font).

| Figure/Supplemantary Figure/<br>Supplementary Table | Medium | Vanco.<br>100 µg/ml | Agar | Culture |
|---|---|---|---|---|
| 1A-D; 1F | BCDA | + | + | regenerated from protoplasts, 5-day-old |
| 1E; 1G | BCD | + | + | regenerated from protoplasts, 5-week-old |
| 2A-D; 2G | BCDA | + | + | regenerated from protoplasts, 5-day-old |
| 2E-F; 2H-I | BCD | + | + | regenerated from protoplasts, 5-week-old |
| 3A-B | BCDA | + | + | regenerated from homogenized protonemata, 1-week-old |
| 3C; 3E, 5 days | BCDA | + | + | regenerated from protoplasts, 5-day-old |
| 3D; 3E, 4 weeks | BCD | + | + | regenerated from protoplasts, 4-week-old |
| 4A-C | BCDA | + | + | regenerated from homogenized protonemata, 1-week-old |
| 4D, 5 days; 4E | BCDA | + | + | regenerated from protoplasts, 5-day-old |
| 4D, 5 weeks | BCD | + | + | regenerated from protoplasts, 5-week-old |
| 4F-G | BCDA | + | + | regenerated from protoplasts, 4-day-old |
| 5A, 5 days; 5C-E, 5 days | BCDA | + | + | regenerated from protoplasts, 5-day-old |
| 5A, 4 weeks; 5E, 4 weeks<br>genotype: WT, $rop^{4xKO}$/$ind^{pro}$:*ROP2* →<br>≤ $2x10^{-5}$ µM β-estradiol;<br>≥ $4x10^{-4}$ µM β-estradiol | BCD | + | + | regenerated from protoplasts, 4-week-old |
| 5A, 4 weeks, inset<br>genotype: $rop^{4xKO}$/$ind^{pro}$:*ROP2* →<br>$2x10^{-4}$ µM β-estradiol | BCD | + | + | regenerated from protoplasts, 4-week-old |
| | | + | - | regenerated from. protoplasts, 4-week-old |
| 5B | BCDA | + | + | regenerated from protoplasts, 7 to 10-day-old |
| 6A | BCDA | + | + | regenerated from homogenized protonemata, 1-week-old |
| 6B-F; 6H 5 days | BCDA | + | + | regenerated from protoplasts, 5-day-old |
| 6G; 6H, 4 weeks | BCD | + | + | regenerated from protoplasts, 4-week-old |
| 7A | BCDA | + | + | regenerated from homogenized protonemata, 1-week-old |
| 7B, 5 days; 7D, 5 days; 7E<br>genotypes: WT; *rop3/2/4;*<br>$rop^{4xKO}$/$ROP1^{pro}$:$ROP1^{F31L}$ | BCDA | + | + | regenerated from protoplasts, 5-day-old |
| 7B, 5 days<br>genotypes: $rop^{4xKO}$;<br>$rop^{4xKO}$/$ROP1^{pro}$:$ROP1^{Q64L}$ | BCDA | + | + | regenerated from homogenized protonemata, 1-week-old |
| 7B, 5 weeks; 7D, 5 weeks<br>genotypes: WT; *rop3/2/4;*<br>$rop^{4xKO}$/$ROP1^{pro}$:$ROP1^{F31L}$ | BCD | + | + | regenerated from protoplasts, 5-week-old |
| 7B, 5 weeks<br>genotypes: $rop^{4xKO}$;<br>$rop^{4xKO}$/$ROP1^{pro}$:$ROP1^{Q64L}$ | BCD | + | + | regenerated from homogenized protonemata, 5-week-old |
| 7C<br>genotypes: WT; *rop3/2/4;*<br>$rop^{4xKO}$/$ROP1^{pro}$:$ROP1^{F31L}$ | BCD | + | + | regenerated from protoplasts, 5-week-old |
| 7C<br>genotype:<br>rop4xKO/ROP1pro:ROP1Q64L | BCDA | + | - | regenerated from homogenized protonemata, 5-week-old |
| 8C | BCD | + | + | regenerated from homogenized protonemata, 5-week-old |
| S1 | BCDA | + | + | regenerated from homogenized protonemata, 1-week-old |
| S5A-D; S5F, 5 days | BCDA | + | + | regenerated from protoplasts, 5-day-old |
| S5E; S5F, 5 weeks | BCD | + | + | regenerated from protoplasts, 5-week-old |
| S6 | BCDA | + | + | regenerated from homogenized protonemata, 1-week-old |

| Figure/Supplemantary Figure/ Supplementary Table | Medium | Vanco. 100 µg/ml | Agar | Culture |
|---|---|---|---|---|
| S7A | BCDA | + | + | regenerated from homogenized protonemata, 1-week-old |
| S7B | BCDA | + | + | regenerated from protoplasts, 4-day-old |
| S7C-E | BCDA | + | + | regenerated from protoplasts, 5-day-old |
| S7F | BCD | + | + | regenerated from protoplasts, 4-week-old |
| S8<br>genotype: WT | BCD | + | - | regenerated from homogenized protonemata, 5-week-old |
| S8<br>genotype: $rop^{4xKO}/ROP1^{pro}:ROP1^{Q64L}$ | BCD | + | - | regenerated from homogenized protonemata, 5-week-old |
| Table S1<br>≤ $9x10^{-4}$ µM β-estradiol | BCDA | + | + | regenerated from protoplasts, 5-day-old |
